# Biogenesis of circular RNAs *in vitro* and *in vivo* from the *Drosophila Nk2.1*/*scarecrow* gene

**DOI:** 10.1101/2024.02.26.582126

**Authors:** Suhyeon Son, Hyunjin Jeong, Gyunghee Lee, Jae H. Park, Siuk Yoo

## Abstract

*scarecrow* (*scro*) encodes a fly homolog of mammalian *Nkx2.1* that is vital for early fly development as well as for optic lobe development. Interestingly, *scro* was reported to produce a circular RNA (circRNA). In this study, we identified 12 different *scro* circRNAs, which are either mono- or multi-exonic forms. The most abundant forms are circE2 carrying the second exon only and bi-exonic circE3-E4. Levels of circE2 show an age-dependent increase in adult heads, supporting a general trend of high accumulation of circRNAs in aged fly brains. Aligning sequences of introns flanking exons uncovered two pairs of intronic complementary sequences (ICSs); one pair residing in introns 1 and 2 and the other in introns 2 and 4. The first pair was demonstrated to be essential for the circE2 production in cell-based assays; furthermore, deletion of the region including potential ICS components in the intron-2 reduced *in vivo* production of circE2 and circE3-E4 by 80%, indicating them to be essential for the biogenesis of these isoforms. Besides the ICS, the intron regions immediately abutting exons seemed to be responsible for a basal level of circRNA formation. Moreover, the replacement of *scro*-ICS with those derived from *laccase2* was comparably effective in *scro*-circRNA production, buttressing the importance of the hairpin-loop structure formed by ICS for the biogenesis of circRNA. Lastly, overexpressed *scro* affected outcomes of both linear and circular RNAs from the endogenous *scro* locus, suggesting that Scro plays a direct or indirect role in regulating expression levels of either or both forms.

## INTRODUCTION

The circRNAs were first discovered in plant viroids and later in almost all organisms from archaea to humans, signifying them as common transcriptional products as linear RNA forms (lineRNAs) (Sanger *et al*. 1976; Memczak *et al*. 2013; Ashwal-Fluss *et al*. 2014; Westholm *et al*. 2014). Interestingly, an abundance of circRNAs varies between tissues, and their expression patterns do not necessarily match with host mRNAs, leading us to speculate that the molecular mechanism governing the biogenesis of circRNAs is divergently evolved from that of lineRNAs (Salzman *et al*. 2013).

The circular structure is formed through a covalent bond between a 5′ splice donor and an upstream 3′ splice acceptor, an event known as ‘back-splicing’ (Jeck *et al*. 2013; Jeck and Sharpless 2014; Wang *et al*. 2014; Li *et al*. 2018; Kristensen *et al*. 2019; Patop *et al*. 2019). Such a non-conventional splicing event is shown to be facilitated by a hairpin structure formed by base-pairing between so-called ‘intronic complementary sequences (ICSs)’ found in the introns flanking the circRNA-coding exon(s) (Dubin *et al*. 1995; Ashwal-Fluss *et al*. 2014; Liang and Wilusz, 2014; Zhang *et al*. 2014; Ivanov *et al*. 2015; Kramer *et al*. 2015; Starke *et al*. 2015; Aktas *et al*. 2017; Knupp *et al*. 2021). No conserved sequences are emerging in the ICSs, implying the uniqueness of individual ICSs and likely an absence of universal *trans*-acting factor(s) binding to ICSs.

Functionally circRNAs add another layer of complexity to gene regulation. One of the known roles of circRNAs is to regulate gene expression as they bind to microRNAs, a function referred to as ‘miRNA sponge’ (Hansen *et al*. 2013; Memczak *et al*. 2013; Zheng *et al*. 2016). Some of them are shown to act as a protein sponge by binding to RNA-binding proteins, while others contain short open-reading frames producing truncated proteins (reviewed in Kristensen *et al*. 2019). However, the biological roles of the majority of circRNAs await further investigations.

Genome-wide deep RNA sequencing has identified circRNAs from numerous genes in various cell and tissue types in *Drosophila* (Westholm *et al*. 2014). Notably, many circRNAs are derived from the brain tissue, suggesting nervous tissue-specific mechanisms regulating circularization and their certain role(s) in this tissue type (Ashwal-Fluss *et al*. 2014; Westholm *et al*. 2014). Interestingly, the circRNAs in the adult brain accumulate with aging perhaps because of the stable nature of the circRNAs (Jeck and Sharpless 2014; Westholm *et al*. 2014; Barrett et al. 2016). A recent study reported a circRNA derived from a *sulfateless* gene (circSfl) is upregulated in insulin mutants, and overexpression of the circSfl extends the lifespan of fruit flies, suggesting a role of accumulated circSfl in longevity (Weigelt *et al*. 2020). Therefore, accumulated circRNAs in aged brains might be associated with the changes in neural structure and/or function during aging. These findings have shown that *Drosophila* is an excellent model system for understanding biogenesis and *in vivo* roles of circRNAs in a gene-specific manner.

The *scro* gene belongs to the NK-2 homeobox family, most members of which are well known to act in regional or cell-type specification in *Drosophila*. *scro* expression is detected predominantly in the central nervous system (CNS) and pharynx (Zaffran *et al*. 2000; Yoo *et al*. 2020). *scro-*null mutations display lethality between late embryonic and early larval stages, indicating that this gene plays a vital function (Yoo *et al*. 2020). Broad expression of *scro* in the central brain, ventral nerve cord (VNC), and optic lobe in larval and adult stages suggest various neuronal roles in both developing and developed CNS (Yoo *et al*. 2020). One such role is to specify neuroblast identity during the optic lobe development as one of the temporal transcription factors (Wang *et al*. 2011; Konstantinides *et al*. 2022). Another known function is to specify the intermediate neural progenitors in the lineage of the dorsal-medial type-II neural stem cells, which eventually give rise to the neurons in the adult central complex (Tang *et al*. 2022). *scro* expression is also detected in a subset of dopaminergic neurons as well as many other neuronal groups in the adult central brain and VNC, suggesting this gene’s likely functioning in differentiated neurons (Yoo *et al*. 2020). It is likely to act as a negative transcription regulator, as transgenic expression of *scro* in the *Drosophila* pacemaker downregulates the *Pigment-dispersing factor* (*Pdf*) gene, an important clock-downstream factor controlling the circadian locomotor activity rhythms (Renn *et al*. 1999; Park *et al*. 2000; Nair *et al*. 2020).

The *scro* gene produces four linear transcript isoforms via alternative splicing (Yoo *et al*. 2020). In this study, we identified 12 different *scro* circRNAs and characterized their temporal expression patterns, showing that the most abundantly expressed forms are circE2 and circE3-E4. We further identified and dissected ICSs essential for the biogenesis of circE2 and circE3-E4 *in vivo* and *in vitro* and addressed key features of the ICS that determine the outcome of circRNAs.

## MATERIALS AND METHODS

### Fly strains

Oregon-R (OR) was used as a wild-type control. The following transgenic lines were used: knock-in *scro* lines, *scro^ΔE2-EGFP^*, *scro^ΔE2-Gal4^*, *scro^ΔE3-EGFP^*, and *scro^ΔE3-Gal4^*, each of which replaces exon-2 or exon-3 with either *EGFP* or *Gal4* coding sequences (Yoo *et al*. 2020); *UAS-scro^HA^*(Nair *et al*. 2020); *hs-Cre* lines (BDSC# 1092 and 34516); *nos-Cas9* line (KDRC# 233). Flies were raised at 25°C in food vials containing 0.85% agarose, 3.75% sucrose, 3% yeast, 8.4% corn meal, 2% Tegosept (a.k.a. methylparaben), and 1% molasses.

### RNA isolation and cDNA synthesis

To analyze the expression of *scro* circRNAs during development, total RNA was purified from various stages of wild-type using the RNeasy system (Qiagen) according to the manufacturer’s protocol with minor modifications. Briefly, the samples were dissolved in 350 µL of RLT lysis buffer with 7 µL of 2 M dithiothreitol (DTT). Following centrifugation, the supernatant was mixed with 70% ethanol and transferred to the spin column. After washing the column with RW1 buffer and RPE buffer, RNAs were eluted with 30 µL RNase-free water. The concentration and purity of the RNAs were measured using MaestroNano^®^ Spectrophotometer (MaestroGen, MN-913). One µg of total RNA was reverse-transcribed by using the ImProm-Ⅱ^TM^ Reverse Transcriptase (Promega). Three types of primers were used for the reverse transcription reaction: oligo d(T) for linear RNA, and either a gene-specific primer (GSP) or random hexamer for both linear and circular RNA. The reaction was performed at 25°C for 5 min, followed by at 42°C for 60 min, and then terminated at 70°C for 15 min.

### RNase R treatment and reverse transcription (RT)-PCR

Total RNAs were treated with exoribonuclease R (RNase R, Epicentre, RNR07250) to destruct lineRNAs. One µg of RNA was incubated for 30 min at 37°C with three units of RNase R or mock-treated by adding distilled water, and then cDNA was synthesized by using a random hexamer as described above. To identify various types of circRNAs, PCR was performed as follows; a 20 µL reaction contained 2 µL of cDNA, 5 µL of GoTaq G2 Green Master Mix (Promega), and 200 µM of primers. The thermal cycling conditions were; 95°C for 3 min (initial denaturation), followed by 40 cycles of 95°C for 1 min (denaturation), 60°C for 1 min (annealing), and 72°C for 1 min (extension), and then by 1 cycle of 72°C for 5 min (final extension).

### DNA constructs for testing ICS

For the transfection assay, a 905-bp fragment containing E2 and flanking intronic sequences was amplified by PCR using the wild-type genomic DNA as a template and scro-I1-NoICS-F and scro-I2-NoICS-R primer set. The PCR product was cloned into *Bam*HI and *Kpn*I sites in the pPacPL vector (*Drosophila* Genomics Resource Center), generating the pNoICS backbone (Fig. 4Aa). To investigate the role of putative ICSs in circRNA production, a 179-bp within intron-1 and a 565-bp fragment within intron-2 were amplified by PCR using primer sets of scro-I1-BF/scro-I1-BR and scro-I2-KF/scro-I2-KR, respectively. These fragments were cloned into *Bam*HI and *Kpn*I sites in pNoICS. According to the orientations of these ICS fragments, we obtained four constructs, pS-ICS-FF, pS-ICS-FR, pS-ICS-RF, and pS-ICS-RR, as illustrated in Fig. 4Ab.

To test the effect of ICS originating from a different gene on circE2 expression, the *scro* ICSs were replaced by those from *laccase2* (Kramer *et al*. 2015). To do this, a 242-bp fragment within intron-1 and a 392-bp one within intron-2 were amplified by PCR using laccase2-I1-BF/laccase2-I1-BR and laccase2-I2-KF/laccase2-I2-KR primer sets, respectively, and then cloned into *Bam*HI and *Kpn*I site in the pNoICS. Four DNA constructs, pL-ICS-FF, pL-ICS-FR, pL-ICS-RF, and pL-ICS-RR, were shown in Fig. 4Ac.

To assess the effect of ICS lengths on the efficiency of *scro* circRNA formation, serial deletions of 21-bp from either 5′ or 3′ end of the 105-bp of ICS in intron-2 were made by employing a fusion-PCR strategy (Cha-Aim *et al*. 2012). For instance, to delete a 21-bp from the 5′ end, the first two PCRs were performed using pS-ICS-FF as a template, and two primer sets (scro-I2-KF/scro-I2-5′-Δa-R and scro-I2-5′-Δa-F/scro-I2-KR). Equimolar amounts of the two overlapping PCR products were mixed and used as a PCR template along with scro-I2-KF and scro-I2-KR primers. The resulting product was cloned into the *Kpn*I site of the pS-ICS-F plasmid carrying only S-ICS1 to generate p5′-Δa construct (Fig. 5A). Similar approaches were used for other deletion constructs. All primers are listed in Supplementary Table 1.

### Quantitative real-time PCR

To quantify the expression levels of lineRNAs and circRNAs of *scro*, cDNA was synthesized with random hexamer as described above. Each PCR contained 1 µL of cDNA, 200 µM each forward and reverse primers, and 10 µL of power SYBR green PCR master Mix (Promega). The real-time qPCR was performed using a fluorescent quantitative detection system (FQD-96A, Bioer). The cycling conditions were 95°C for 2 min, followed by 40 cycles of 95°C for 15 sec and 60°C for 1 min. Either *Rp49* (*ribosomal protein 49*) or *act5C* (*actin 5C*) was used as a control for normalization. The relative expression levels of RNAs were evaluated by the ΔΔCt -based method using LineGene 9600 Plus software (Bioer) and showed the log10-fold difference in Fig. 3B and Fig.4B. The experiments were repeated at least three times, and the results were presented as mean ± standard deviation (std).

PCR primers are shown in Supplementary Tables 2 and 3.

### Cell culture and transfection assay

Schneider’s 2 (S2) cells were cultured in Schneider’s Insect Medium (WELGENE, South Korea) supplemented with 10% fetal bovine serum (Cytiva, Austria) and 1% penicillin/streptomycin (Cytiva, Austria). The cells were maintained in an incubator at 23°C and subcultured at 80-90% confluency (about once every week). To assess the effect of ICS on the formation of circRNAs, S2 cells were cultured in a 6-well plate to 60% confluence, transfected with 4 µg of plasmid DNA using the FuGENE^®^ HD Transfection (Promega) for 24 h, and then total RNA was isolated from the transfected cells for RT-qPCR.

### Generation of a *scro* mutation lacking ICS

To identify candidate ICSs within the *scro* introns, all intron sequences annotated in the Flybase (https://flybase.org/) were aligned using EMBOSS (http://emboss.toulouse.inra.fr/cgi-bin/emboss/einverted) with the following parameters: ICS for circE2 with the Gap penalty=40, the min score threshold=150, the match score=300, mismatch score=30, and the max extent of repeats=250; ICS for circE3-E4 with the Gap penalty=5, the min score threshold=0, the match score=150, mismatch score=-3, and the max extent of repeats=150; ICS in *laccase2* with the Gap penalty=5, the min score threshold=250, the match score=169, mismatch score=20, and the max extent of repeats=475 options (Supplementary Fig. 2).

To delete the 1180-bp region containing ICS from the intron-2, we employed a CRISPR/Cas9-mediated genome editing system. The target cleavage sites were selected using the flyCRISPR Optimal Target Finder tool (Gratz *et al*. 2014; http://tools.flycrispr.molbio.wisc.edu/targetFinder/). Two complementary sets of gRNA oligonucleotides (Supplementary Table 1), ΔI2-ICS-gRNA1-S/ΔI2-ICS-gRNA1-AS and ΔI2-ICS-gRNA2-S/ΔI2-ICS-gRNA2-AS were annealed, cloned into the pU6-Bbs-chiRNA at *Bbs*I site, and then injected into *nos-Cas9* embryos. Approximately 50 G0 flies were individually crossed with *y w*, and 10 G1 flies from each G0 line were singly mated with *y w*; *Sb*/*TM6B* balancer stock. After 4 days, G1 flies were screened by single-fly PCR using a primer set I2F/I2R (Supplementary Table 2); the deletion line (ΔI2-ICS) is expected to produce a 245-bp PCR product, instead of the 1552-bp fragment from the wild-type. Since the positive G1 flies are heterozygous for the ΔI2-ICS allele, positive G2/TM6B flies were crossed to the balancer to establish the stocks (Supplementary Fig. 3). As a result, we established two lines, namely *ΔI2-ICS-44* and *ΔI2-ICS-45*, which carry the identical genomic lesion confirmed by sequencing.

## RESULTS

### Verification of circRNAs from *scro* transcripts

The *scro* is annotated to produce four linear transcripts (*RA*, *RB*, *RC*, and *RF*) via alternative splicing (Fig. 1A). It was also reported that a circRNA carrying a single exon originated from this locus (Westholm *et al*. 2014). To see if there are additional *scro* circRNA isoforms, total RNAs were extracted from wild-type adult heads where their linear transcripts are most abundantly expressed (Yoo *et al*. 2020). We designed a set of convergent primers (E2F2 and E2R2) that are intended to detect both linear and circular forms, and a set of divergent primers (E2F1 and E2R2) that should detect only circular forms (Fig. 1B). When we tested oligo d(T)-primed cDNA samples that derived from lineRNAs, convergent primers generated PCR product (Fig. 1C, lane d(T) in upper panel) but divergent primers did not (Fig. 1C, lane d(T) in lower panel). However, with hexamer-primed cDNA samples, PCR products were generated by both convergent and divergent primers (Fig. 1C), indicating the presence of a circRNA from E2.

**Fig. 1.**
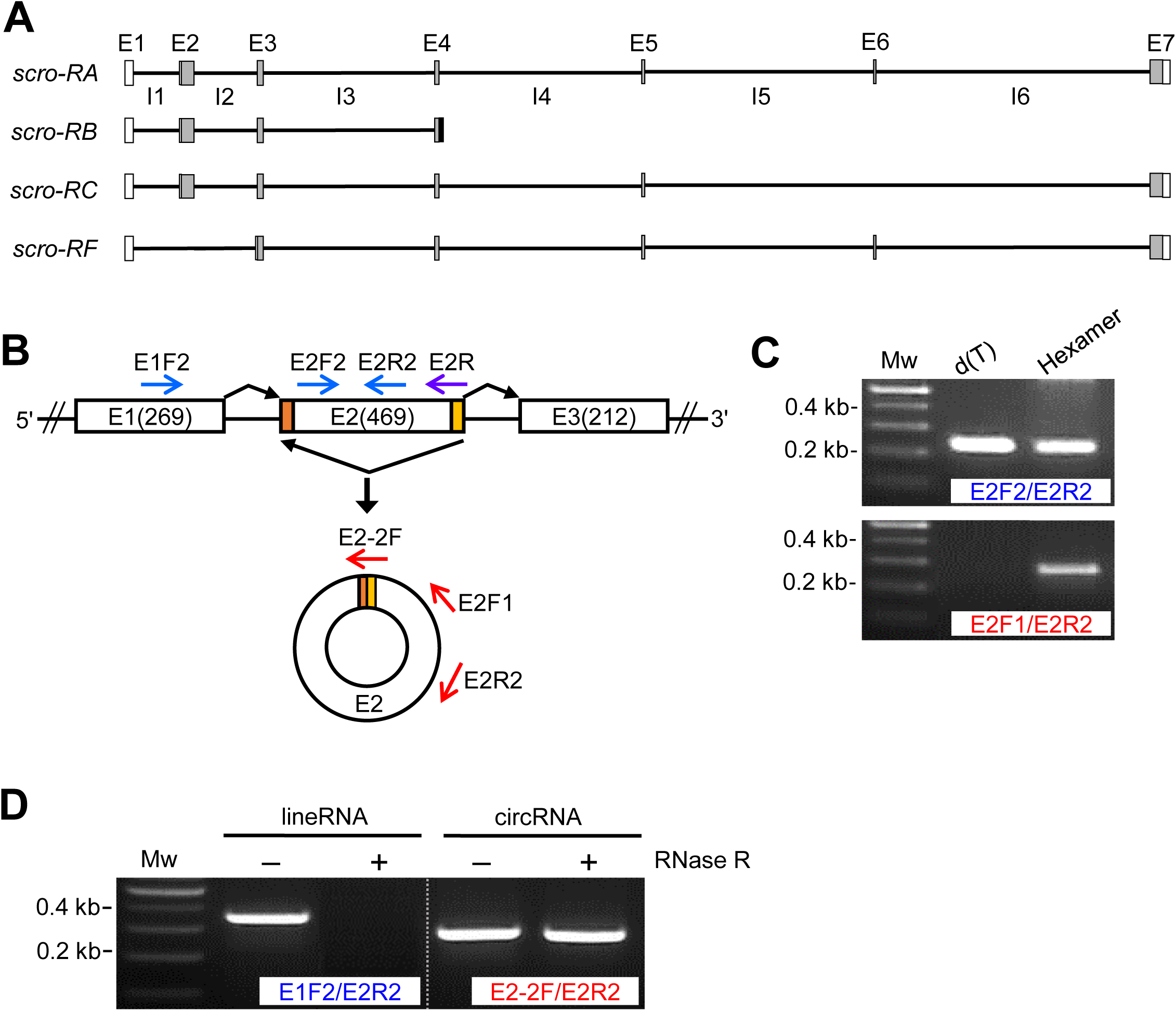
Verification of circRNA derived from the *scro* locus. **(A)** Exon (E)-intron (I) genomic organization of *Drosophila scro* containing eight exons (E1∼E7 and *RB*-specific exon). According to FlyBase, the four mRNA isoforms are generated by alternative splicing. White boxes, UTRs; gray boxes, coding regions; black box, *RB*-specific exonic sequence, otherwise, it is an intronic region for the others. **(B)** Schematic diagram of RNA splicing leading to the generation of *scro* circE2. The noncanonical exon junction point is formed when the 5′ splice site of E2 (yellow) is joined to the 3′ splice site at the beginning of E2 (orange) via back-splicing. E2R primer (violet arrow) was used for cDNA synthesis, and the convergent primers (blue arrows) and divergent primers (red arrows) were used for PCR. **(C)** RT-PCR results using E2F2/E2R2 primer set (upper panel) and E2F1/E2R2 primer set (lower panel). The primers used for cDNA synthesis are shown above each lane. PCR fragments were amplified using outward primers in cDNA samples synthesized from random hexamer, but not in cDNAs from oligo d(T) primer (lower panel), indicating the presence of circE2. Mw is a 100-bp DNA ladder. **(D)** Effect of RNase R treatment on the detection of circRNA. Total RNA samples were either mock-treated (**‒**) or RNase R-treated (+) before the cDNA synthesis using hexamer. In RNase R-treated RNA samples, only cDNAs using exon junction primer (E2-2F) were amplified, verifying the existence of *scro*-circRNAs. Mw represents a 100-bp DNA ladder.

**Fig. 2.**
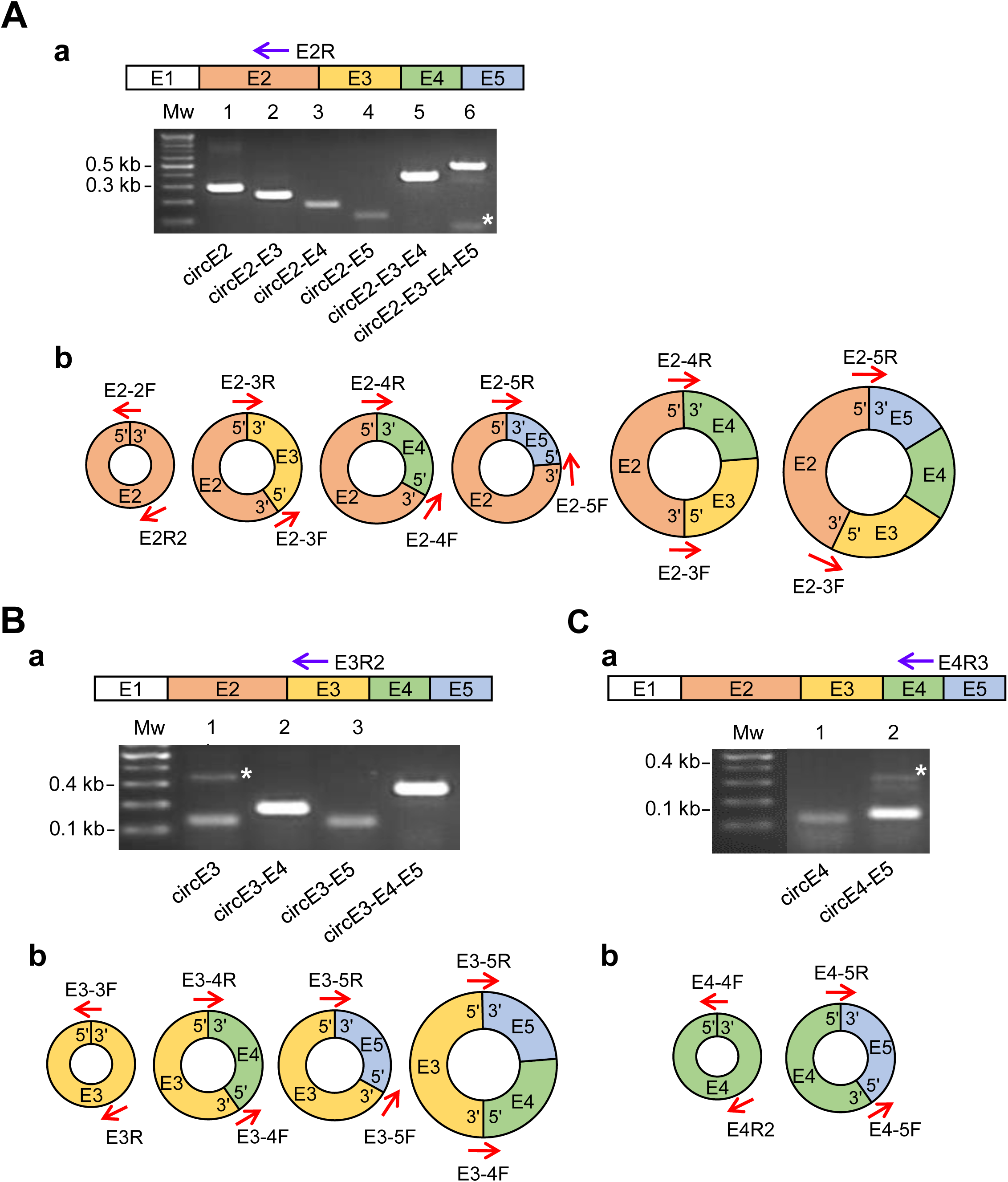
Identification of different types of *scro* circular RNAs. To detect various types of exonic circRNAs, cDNAs were synthesized with gene-specific primers (violet arrows above agarose gels), and RT-PCR was performed using the junction primers spanning the exon-exon joining regions (red arrows in diagrams). In gel images, non-specific bands are indicated by asterisks, and Mw represents a 100-bp DNA ladder. Schematic diagrams represent *scro*-circRNAs carrying one to four exons. Each exon is differently color-coded. **(A)** Six types of circRNAs detected from E2R-primed cDNA samples are shown in agarose gel **(a)** and diagrams **(b)**. The circRNA types corresponding to each lane are indicated at the bottom of the agarose gel. The sizes of PCR fragments are as follows; lane 1, 297-bp; lane 2, 242-bp; lane 3, 182-bp; lane 4, 126-bp; lane 5, 394-bp; lane 6, 490-bp. **(B)** Four types of circRNAs detected from E3R2-primed cDNA samples are shown in agarose gel **(a)** and diagrams **(b)**. Lane 1, 146-bp; Lane 2, 178-bp; Lane 3, 122-bp; Lane 4, 274-bp. **(C)** Two types of circRNAs detected from E4R3-primed cDNA samples are shown in agarose gel **(a)** and diagrams **(b)**. Lane 1, 122-bp; Lane 2, 123-bp.

To ascertain the foregoing result, total RNAs were treated with RNase R to digest linear RNAs before the cDNA synthesis, and then PCR was carried out using either convergent primers (E1F2 and E2R2) for lineRNAs or junction and divergent primers (E2-2F and E2R2) for circRNAs (Fig. 1B). As expected, lineRNA-targeting primers produced PCR product from mock-treated RNA, but none from the RNase R-treated one (Fig. 1D, left panel). In contrast, circRNA-targeting primers produced PCR products from both RNA samples (Fig. 1D, right panel). The results validate our approach being effective in finding exonic *scro* circRNAs.

### Identification of 12 different circRNA forms from *scro*

To explore additional E2-containing circRNAs, we designed six different sets of primers as illustrated, each of which was intended to detect a specific combination of E2-containing multi-exonic circRNAs (Fig. 2Ab). As a result, we identified six distinct PCR products using cDNAs generated from RNase R-treated total RNAs, which are referred to as circE2, circE2-E3, circE2-E4, circE2-E5, circE2-E3-E4, and circE2-E3-E4-E5, respectively (Fig. 2Aa). The sequencing of these products showed the lack of intronic region, verifying characteristic back-splicing events for all identified circRNAs (Supplementary Fig. 1A). By extending this strategy to other exons, we identified four additional circRNAs containing the E3 (circE3, circE3-E4, circE3-E5, and circE3-E4-E5) (Fig. 2Ba-b, Supplementary Fig. 1B), and two including the E4, circE4 and circE4-E5 (Fig. 2Ca-b, Supplementary Fig. 1C). Despite our painstaking efforts, we were unable to find circRNAs including E1, E6, and E7. It is consistent with previous findings showing that the first or last exon of a gene is rarely made into circRNAs, presumably because they are flanked by only one intron (Salzman *et al*. 2012; Lasda and Parker 2014; Westholm *et al*. 2014; Gruner *et al*. 2016). However, it was surprising not to see circRNAs carrying E6, although this exon is flanked by the two largest introns. The circRNAs can be sorted into three types based on their intronic and/or exonic compositions: exonic circRNAs, intronic circRNAs, and exon-intron circRNAs (Lasda and Parker 2014; Shen *et al*. 2015). All of the 12 identified *scro* circRNAs are either single- or multi-exonic, the latter of which suggests that the inter-exonic introns are precisely excised out.

### Differential expression of *scro* circRNA isoforms

Expression levels of each circRNA were assessed from different developmental stages by quantitative real-time PCR (qPCR). Expression of circE2 and circE3-E4 types was readily detected from embryonic to adult stages (Fig. 3A). In general, circE2 levels are about 30% higher than circE3-E4. On the other hand, other circRNAs were weakly detected in all of the stages examined compared to circE2 and circE3-E4. These results indicate that circE2 and circE3-E4 are the two most prevalent forms expressed throughout development and in the adult stage.

**Fig. 3.**
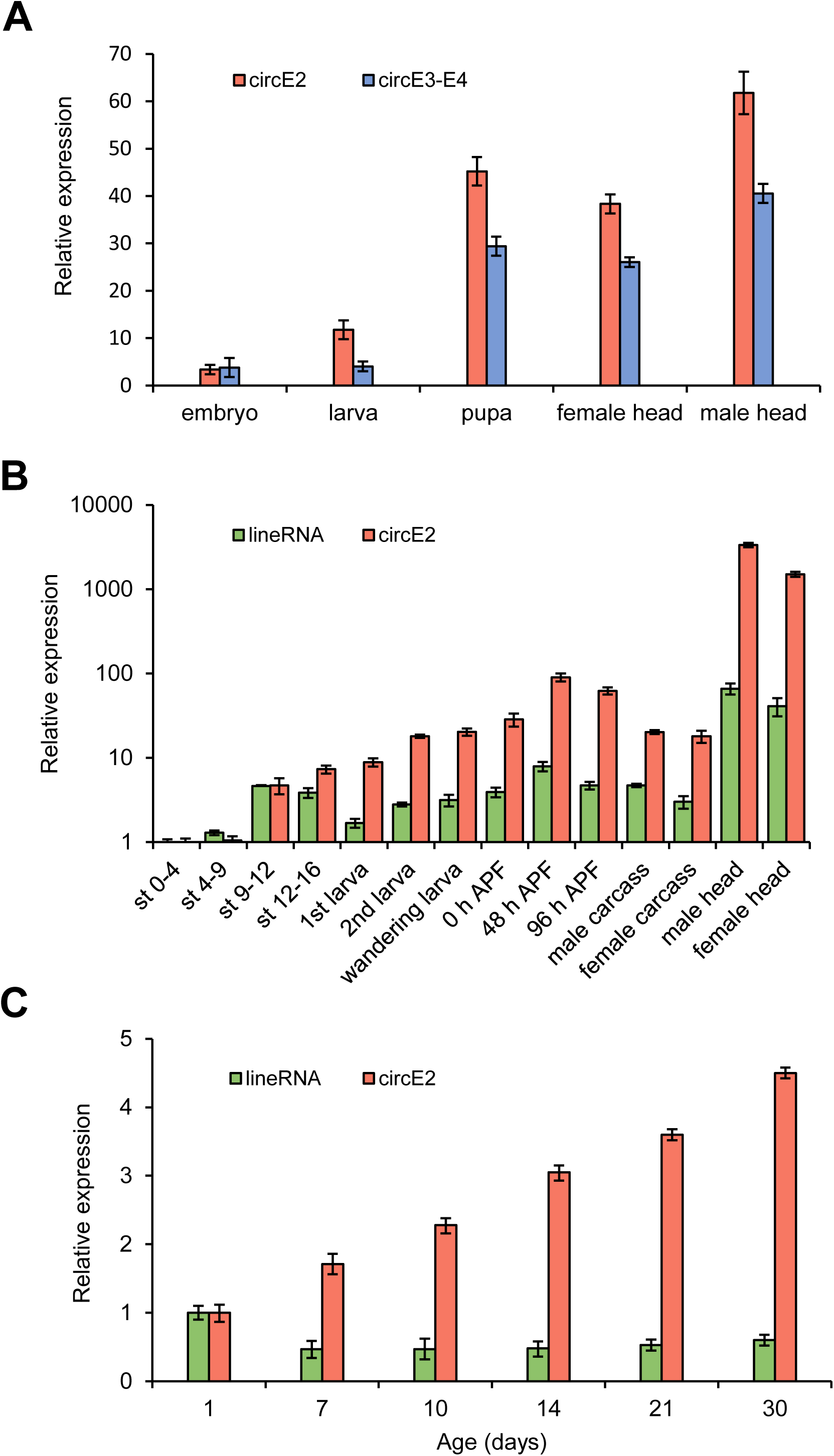
Expression levels of circRNAs during development and aging by real-time PCR. **(A)** Expression levels of the two predominant circRNAs (circE2 and circE3-E4) at major developmental stages. Note that the levels of circE2 are greater than circE3-E4. The levels of other types of circRNAs are too low to measure by real-time PCR (not shown). E2-2F/E2R2 primer set was used for circE2, and E3-4F/E3-4R set for circE3-E4. **(B)** Developmental expression profiles of lineRNAs and circE2. E2-2F/E2R2 primers were used to detect circRNA, and E1F2/E2R2 primers for linear RNAs. APF, after puparium formation. **(C)** Aging-dependent expression levels of lineRNAs and circE2. The circRNA expression levels increase in an age-dependent manner, while lineRNA levels decrease by 50% within 7 days. Each bar represents mean ± standard deviation (std).

Next, we compared the levels of circRNA and lineRNA forms in finely staged animals during development. We focused on the circE2 as it is the major isoform. Levels of circE2 were compared with combined levels of the three linear transcripts containing E2 (*RA*, *RB*, and *RC*; Fig. 1A). For the latter, E1F2 and E2R2 primers were used to cover the region that is common to all three linear isoforms (Fig. 1B). Expression levels of both circular and linear forms were comparable up to embryonic stages 9-12; afterward, circE2 levels increased exponentially above lineRNAs (Fig. 3B).

The highest levels of both forms were found in adult heads, which is consistent with the CNS being the main *scro* expression site (Fig. 3B; Yoo *et al*. 2020). Of interest, most of the *Drosophila* circRNAs detected in the brain were shown to accumulate with age (Westholm *et al*. 2014; Weigelt *et al*. 2020). This prompted us to examine whether circE2 expression also follows this trend.

Indeed, the levels of circE2 increased nearly 4.5-fold over 30 days. Meanwhile, lineRNA levels reduced by half within 7 days of aging and then remained flat afterward (Fig. 3C).

### Intronic sequences required for the biogenesis of *scro* circRNAs

Intronic complementary sequences (ICSs) located in introns flanking a circRNA exon or the first and last exons of multi-exonic ones are known to play a critical role in the production of circRNAs, as they facilitate the formation of the stem-and-loop structure in the pre-mRNA and then back-splicing of the paired introns to release the circRNAs (Ivanov *et al*. 2015; Wang *et al*. 2019).

Potential ICSs were recognized in many of the *Drosophila* genes producing circRNAs (Ashwal-Fluss *et al*. 2014; Kramer *et al*. 2015; Weigelt *et al*. 2020), but their actual involvement in the circularization was validated only in a small number of genes including *muscleblind* (*mbl*) and *laccase2* genes (Ashwal-Fluss *et al*. 2014; Kramer *et al*. 2015). By aligning sequences of introns flanking exons of circE2 and circE3-E4, we found two putative intronic pairs consisting of at least 80-105 nucleotides; one in intron-1 and intron-2 and the other in intron-2 and intron-4, which are possibly involved in the production of circE2 and circE3-E4, respectively (Supplementary Figs. 2A and B).

We wanted to test the first candidate ICSs in cultured cells to see if they are indeed required for the biogenesis of the *scro* circE2. We made a plasmid containing E2 and flanking intronic regions lacking putative ICSs (pNoICS, Fig. 4Aa). Cells transfected with pNoICS produced low levels of circE2, but significantly higher when compared to those with the blank plasmid (mock), indicating that the immediately flanking intronic regions are capable of circularizing E2 albeit at low frequency (Fig. 4B).

**Fig. 4.**
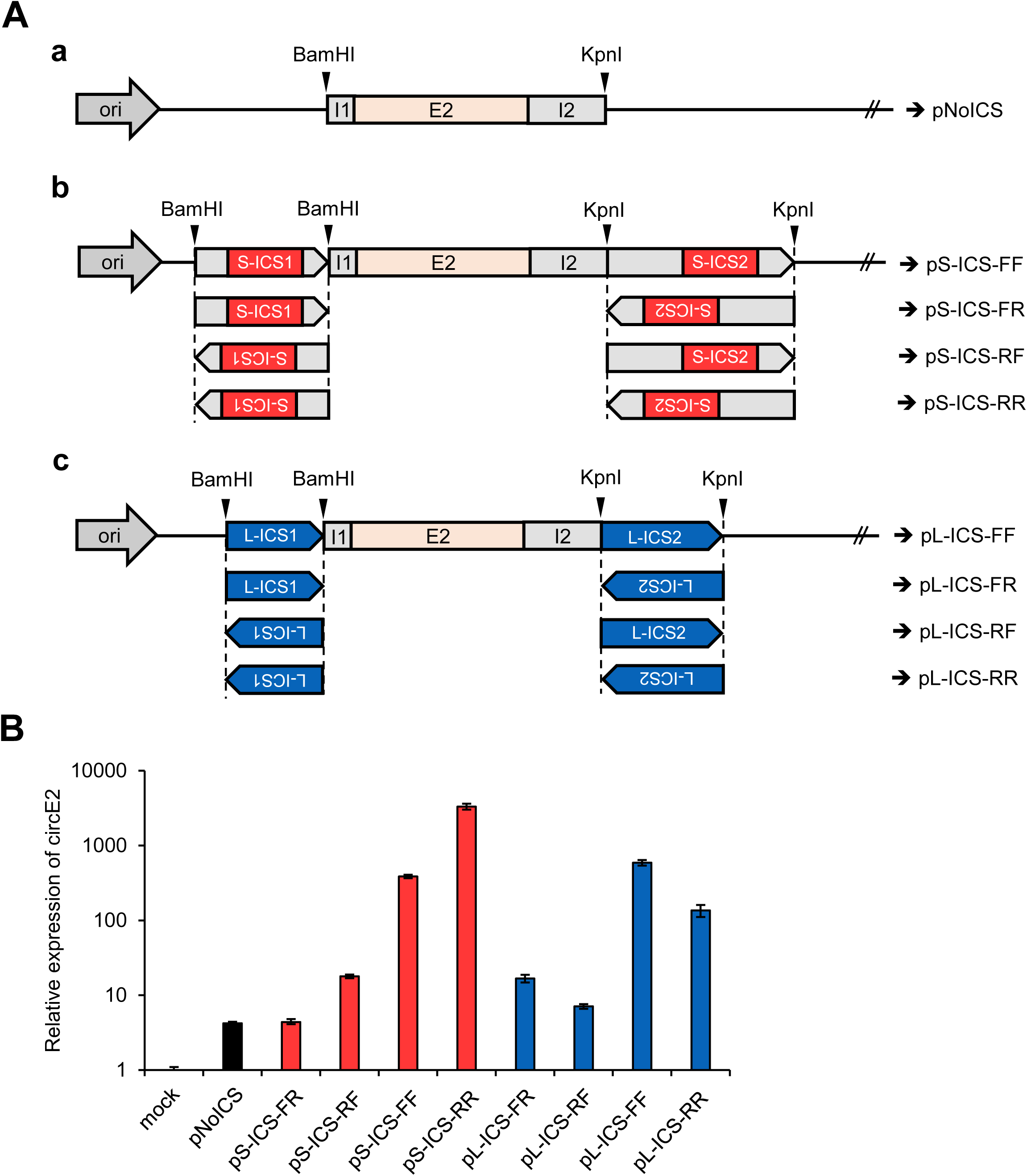
Effects of ICS orientation and origin on circE2 expression *in vitro*. **(A)** Schematic diagrams of DNA constructs for transfection assay. **(a)** pNoICS was generated by cloning of 905-bp PCR fragment carrying *scro* E2 and the flanking intronic sequences into the pPacPL vector. **(b)** The 179-bp containing *scro* intron-1 ICS (S-ICS1) and 565-bp containing *scro* intron-2 ICS (S-ICS2) fragments were cloned into pNoICS at *Bam*HI and *Kpn*I sites, respectively, generating four constructs according to their orientations. The first letter following ICS designates the orientation of intron-1 ICS, and the second letter does intron-2 ICS. ‘F’ stands for the same direction as genomic DNA, and ‘R’ for reverse orientation. **(c)** The 242-bp containing *laccase2* intron-1 ICS (L-ICS1) and 392-bp containing *laccase2* intron-2 ICS (L-ICS2) fragments were cloned into pNoICS at *Bam*HI and *Kpn*I sites, respectively, generating four constructs according to their orientations. **(B)** Real-time PCR results showing circE2 levels in transfected S2 cells. The expression levels of circE2 (red bars) in both pS-ICS-FF and pS-ICS-RR are much higher compared to those in pS-ICS-FR and pS-ICS-RF (S stands for *scro*), strongly implying the importance of hairpin structure for the biogenesis of circRNA. Similar results were obtained when *scro* ICS was replaced with *laccase2* ICS (blue bars) as the hairpin structures are expected to form in both pL-ICS-FF and pL-ICS-RR, but not in pL-ICS-FR and pL-ICS-RF (L stands for *laccase2*), indicating that the origin of ICS is not a crucial factor for circRNA formation. Each bar represents mean ± std.

We consider this basal level expression. Next, extended intron-1 and intron-2 fragments including the 105-bp ICSs were tested in their natural orientations (pS-ICS-FF, shortly FF; ‘S’ represents *scro* and ‘Forward (F)’ indicates the same orientation of ICS as in the *scro* genomic sequence). Remarkably, the FF construct substantially increased circE2 levels, indicating ICS’s essential role in enhancing circE2 production levels (Fig. 4B). These results together strongly support that the hairpin structure formed by the ICS enhances circE2 biogenesis, while intronic regions immediately flanking E2 are necessary for the basal level of circE2 production, thereby supporting that both sites are needed to give rise to full production of circE2.

To further confirm if the base-pairing of the ICSs is essential for the circularization, we tested reversely oriented ICSs (RR) (Fig. 4Ab). Since both FF and RR can similarly make base-pairing in theory, the RR construct would facilitate circE2 production as equally as FF does. Intriguingly, the RR produced the circE2 at even higher levels than FF did (Fig. 4B). We reasoned that it might be due to the closer proximity of S-ICS2 to the E2 in the RR context, which somehow promotes the hairpin formation. If this is the case, the distance between ICSs and circularizing exon could be another factor that determines the efficacy of back-splicing. In contrast to FF and RR, circE2 expression from FR was at the basal level, while that from *RF* was slightly higher than FR (Fig. 4B). Together the data showed that both base-pairing of ICSs and their proximity to the exon determine output levels of circRNAs.

Our next question is whether the formation of hairpin structure mediated by ICS, regardless of its origin, is essential for the biogenesis of *scro*-circRNA. To test this, we employed the ICSs found for a *laccase2* circRNA (Kramer *et al*. 2015; Supplementary Fig. 2C). The *laccase2* ICSs (L-ICS1 and L-ICS2) were inserted into pNoICS to generate pL-ICS-FF, -FR, -RF, and -RR (Fig. 4Ac). Both FF and RR constructs produced significantly higher levels of circE2 than FR or RF did (Fig. 4B). The *laccase2* FF outcome was comparable to that of *scro* FF, while *laccase2* RR was less effective than *scro* RR. Together these results support that ICS-mediated hairpin is a key structural feature for the production of circE2.

### Finding a core region of ICS essential for the productivity of circE2

To further investigate if there is a minimal or core region within the ICS for promoting circE2 production, the 105-nt ICS for circE2 located in intron-2 (S-ICS2) was divided equally into five 21-nucleotide (nt)-long regions labeled from ‘a’ to ‘e’ (Figs. 5A and C). These regions were serially deleted in the 5’ → 3’ direction and designated as p5’-Δa, p5’-Δab, p5’-Δabc, and p5’-Δabcd, respectively (blue bars in Fig. 5A). Each of these constructs was tested in S2 cells for circE2 production. Expression levels of circE2 in p5’-Δa, p5’-Δab, and p5’-Δabc seemed slightly higher or comparable to FF control, whereas p5’-Δabcd produced circE2 at the basal level as the negative control (pΔabcde) did (blue bars, Fig. 5B). These results inform that the ‘d’ region is critical for the hairpin formation. We also reasoned that higher yields of p5’-Δa and p5’-Δab might be because the deletion of these regions brought the ICS closer to E2.

**Fig. 5.**
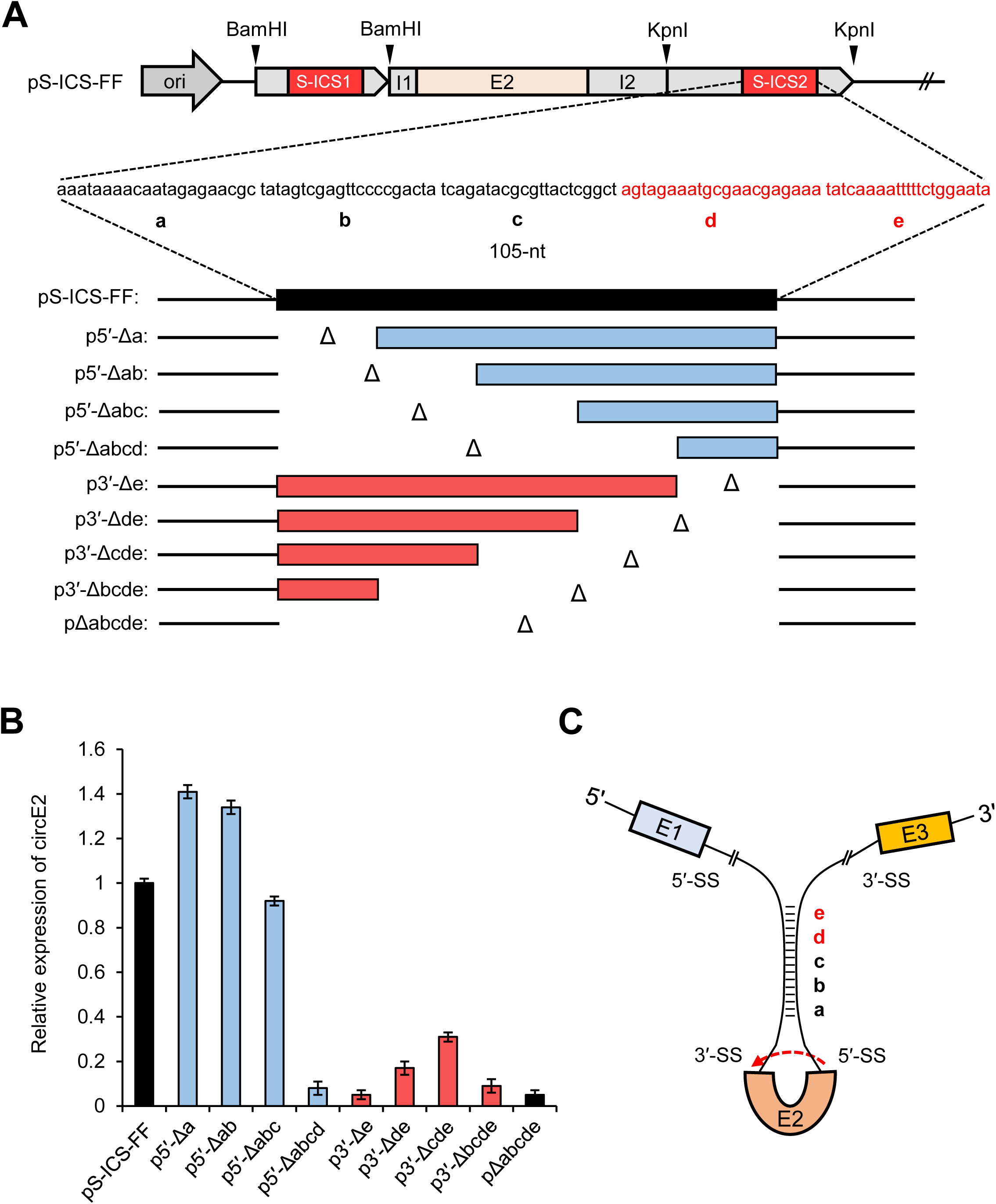
Effect of ICS length on circE2 expression. **(A)** Schematic diagrams of the deletion constructs. The 105-nt of ICS in *scro* intron-2 was serially removed by 21-nt as represented by ‘a’ through ‘e’ from the pS-ICS-FF. **(B)** Real-time PCR results of analyzing the expression levels of circE2 after transfection of each DNA construct into the S2 cell line. Primers used for cDNA synthesis and PCR were hexamer and E2-2F/E2R2, respectively. Comparable circE2 levels between p5′-Δabc and pS-ICS-FF (positive control) suggest that at least a 40-bp length of ICS is required for circE2 formation. However, transfection of p3′-Δde greatly reduced expression in circE2, suggesting that both ‘d’ and ‘e’ regions are critical for circE2 production. Each bar represents mean ± std. **(C)** Schematic of the hairpin loop involved in the biogenesis of circE2. The dashed arrow indicates back-splicing. Distinct regions within ICS are indicated by a-e. ss, splicing site; I, intron.

Next, the serial deletions were made in the 3’ → 5’ direction, generating p3’-Δe, p3’-Δde, p3’-Δcde, and p3’-Δbcde, respectively (red bars in Fig. 5A). All of these constructs showed basal or slightly higher than basal levels of circE2 expression (red bars in Fig. 5B). We were surprised to see the lack of activity by p3’-Δe missing only ‘e’ region, because it includes the ‘d’ that we expected to be essential from the p5’-Δabcd result. Hence, we propose that the sequence spanning both ‘d’ and ‘e’ regions serves as a core ICS motif. The ‘d-e’ region is the most distal part of the stem, positioned away from the loop region (Fig. 5C). It leads us to speculate that the ‘d-e’ region might promote hairpin formation or interact with *trans* factor(s) that play a role in hairpin formation or circularization. To verify if the core ICS is involved in hairpin structure, we aligned these sequences using EMBOSS that found inverted repeats needed for stem-loops in nucleotide sequences. However, except circE2 and circE3-E4, none of the particular nucleotides needed for significant hairpin structure formation was found in other *scro*-circRNAs.

### *In vivo* role of S-ICS2 for the production of *scro* circRNA

To further demonstrate the role of the identified S-ICSs for circE2 expression *in vivo*, we employed the CRISPR/Cas9-mediated genome editing tool to remove an 1180-bp intronic region including the S-ICS2 and obtained two independent lines (Δ*I2-ICS-44* and *-45*) (Fig. 6A and Supplementary Fig. 3). In contrast to early lethality of the *scro*-null mutants (Yoo *et al*. 2020), the new mutant flies showed no recognizable defects in their development and fertility. Real-time qPCR demonstrated that the expression levels of circE2 reduced by about 80% in the adult head of both mutant lines (Fig. 6Ba).

**Fig. 6.**
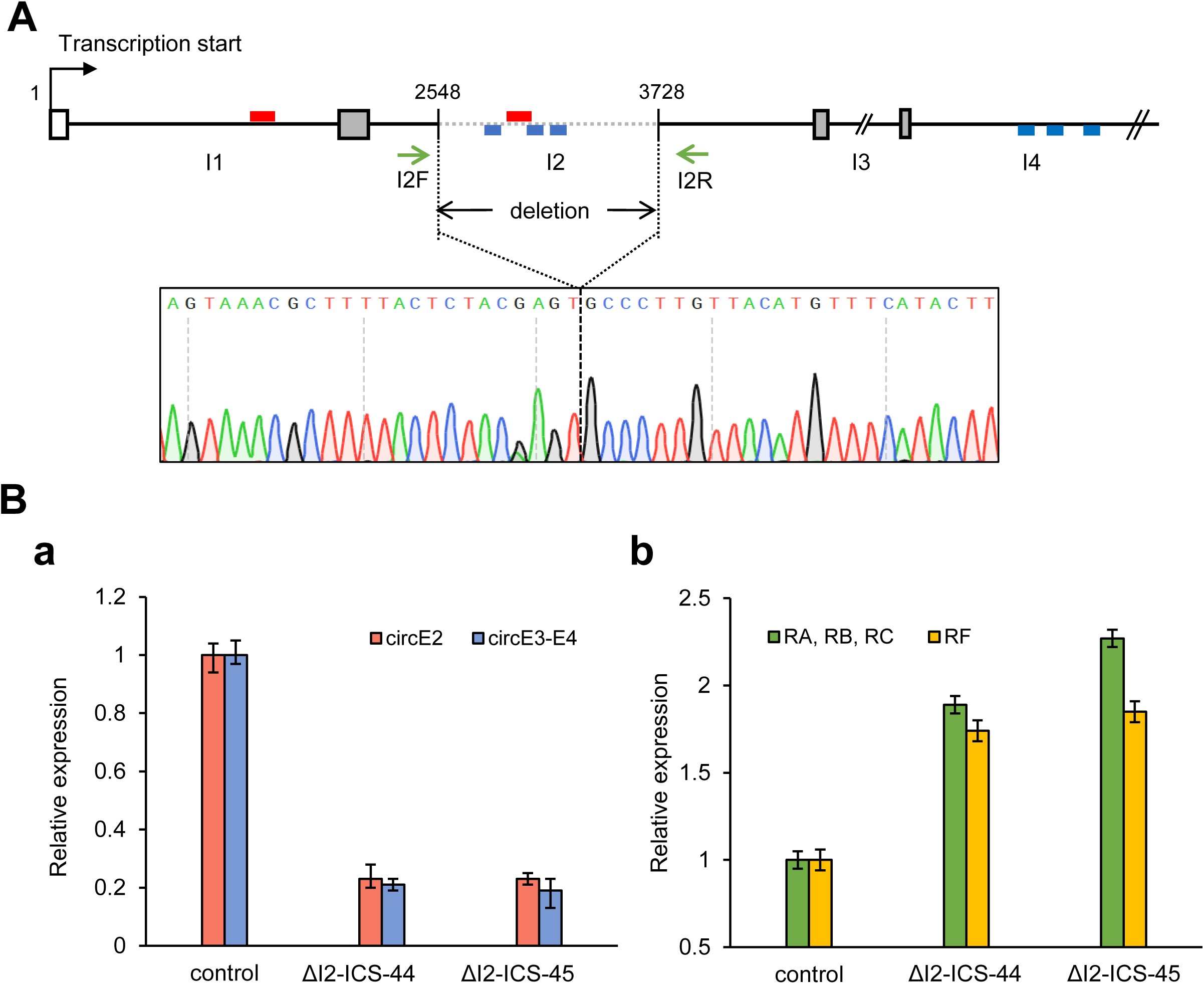
Analysis of the role of ICS *in vivo*. **(A)** Schematic diagram showing the cleavage sites directed by gRNAs. The gRNAs were designed to remove the 1180-bp region containing ICS within the intron-2 (I2). The red boxes represent ICS for circE2, and the blue boxes represent ICS regions for circE3-E4. PCR using I2F and I2R primers was used to determine the deleted region. Both deletion lines carry the same lesion as shown by the sequencing data. **(Ba)** Real-time PCR analysis to determine the expression levels of circRNAs from the two homozygous deletion lines, *ΔI2-ICS-44* and *ΔI2-ICS-45*. Not only circE2 but circE3-E4 levels are dramatically reduced as compared to those of the control. E2-2F and E2R2 primers were used for the detection of circE2, and E3-4F and E3-4R primers for circE3-E4. Bars represent mean ± std. **(Bb)** Real-time PCR result shows the expression levels of linear transcript isoforms in the two homozygous deletion lines, *ΔI2-ICS-44* and *ΔI2-ICS-45*. In contrast with the **(Ba)** result, *scro* lineRNA levels are increased about 2-fold than those of the control. E1F2 and E2R2 primers were used for the detection of linear transcripts containing E2 (*RA*, *RB*, and *RC*) and E1-3F and E3R primers for *RF* transcript. Bars represent mean ± std.

The deleted region contains not only the S-ICS2 essential for circE2 but also another potential ICS for circE3-E4 production as shown in Supplementary Fig. 2B, thus expecting to affect circE3-E4 production. Indeed, circE3-E4 levels showed a significant reduction like circE2 in the mutants (Fig. 6Ba). Taking these data together strongly supports that ICSs are important for the *in vivo* production of both circE2 and circE3-E4. Despite the significant reduction of circRNA levels, since the mutants look quite normal, either low levels of circE2 or circE3-E4 are still sufficient for functioning or they are dispensable for larval and adult development. Complete elimination of the circRNAs will address their *in vivo* roles more clearly. We also like to note that expression levels of two types of lineRNAs, one type including E2 (*RA*, *RB*, and *RC*) and the other E3 (*RF*), were greater by about 2-fold in Δ*I2-ICS-44* and *-45* mutants than in control (Fig. 6Bb). A likely possibility is that conventional splicing for the lineRNAs and back-splicing for the circRNA compete for the same splicing machinery.

### Generation of circRNAs in *scro* knock-in mutants

In addition to ICSs, we wanted to test if there is any exonic contribution to circRNA production. We utilized previously generated *scro* knock-in mutants; *scro^ΔE2-EGFP^*, *scro^ΔE3-EGFP^*, *scro^ΔE2-Gal4^*, and *scro^ΔE3-Gal4^* (Yoo *et al*. 2020). *scro^ΔE2-EGFP^* was made by replacing most E2 with a cassette including EGFP, 3XP3-RFP marker, and SV40 terminator. For *scro^ΔE3-EGFP^*, the E3 and part of the flanking intron-3 were replaced with the same cassette. *scro^ΔE2-Gal4^*and *scro^ΔE3-Gal4^* (Gal4 knock-in) are the same as EGFP knock-in, except for the Gal4 coding sequence in place of EGFP. Since SV40 terminator blocks the process of transcription, we first removed SV40 and 3XP3-RFP by using Cre/lox recombination system (Fig. 7). These excision lines were distinguished from the original lines by adding an asterisk to the end as follow; ‘*scro^ΔE2-EGFP*^* or *scro^ΔE2-Gal4*^*, and ‘*scro^ΔE3-EGFP*^* or scro*^ΔE3-Gal4*^*.

**Fig. 7.**
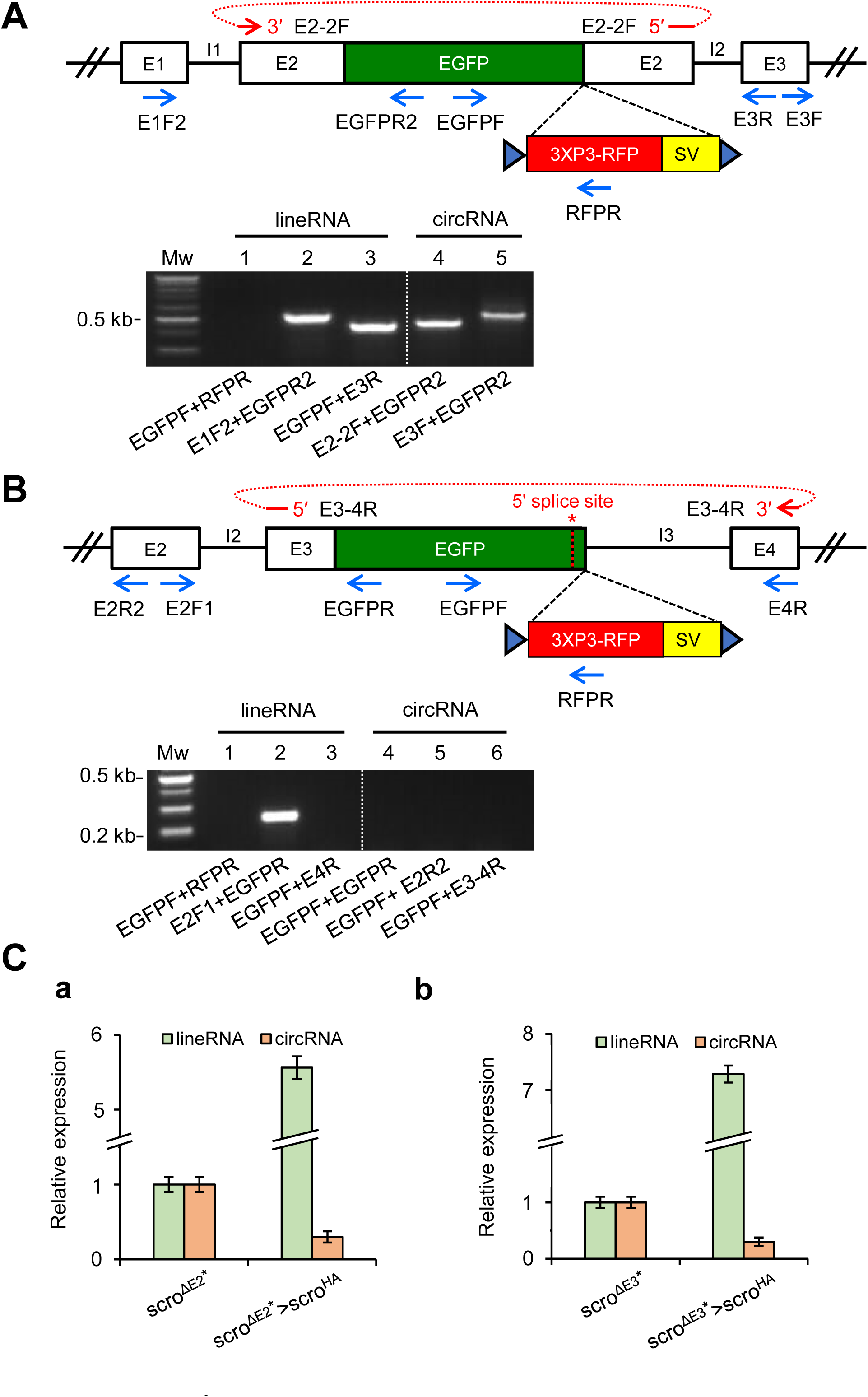
Analysis of circular transcripts from *scro* knock-in mutants. **(A)** Genomic organization of *scro^ΔE2-EGFP*^* knock-in mutant (upper panel). The RFP marker and SV40 termination signal were removed from *scro^ΔE2-EGFP^* by using the Cre/loxP recombination system. The blue triangle represents the loxP sequence, and the red arrow above the diagram represents the E2-2F exon junction primer. RT-PCR was performed by using the primers indicated under the gel image (lower panel). No PCR product in lane 1 verifies the removal of the RFP marker and SV40 terminator. The lineRNAs carrying EGFP are produced normally (lanes, 2 and 3). Note that circRNAs carrying E2/EGFP exon are also formed by back-splicing (lane 4, circE2/EGFP; lane 5, circE2/EGFP-E3. **(B)** Genomic organization of *scro^ΔE3-EGFP*^* knock-in mutant (upper panel) and RT-PCR results (lower panel). The red arrow in the diagram represents the E3-4R exon junction primer. Cre/loxP-mediated elimination of the RFP marker and SV40 terminator was confirmed in lane 1. Analysis of linear transcripts showed that the splicing of intron-2 occurs normally (lane 2), whereas intron-3 is not spliced out (lane 3). Any circRNAs containing EGFP (lane 4), circE2-E3/EGFP (lane 5), and circE3/EGFP-E4 (lane 6) are not produced, most likely due to the lack of the 5′ splice site in the intron-3 (red asterisk above the diagram). **(C)** Real-time qPCR analysis to determine the expression levels of linear and circular RNAs in response to transgenic overexpression of *scro* lineRNA. Either the *scro^ΔE2-Gal4*^* **(a)** or *scro^ΔE3-Gal4*^* line **(b)** was crossed to a *UAS-scro^HA^*, and then RNA samples were prepared from the adult F1 heads. qPCR was performed by using E1F2/E2R2 primer set for lineRNAs and E2-2F/E2R2 primer set for circE2. The expression of lineRNAs increased more than 5-fold, while circE2 levels reduced less than 3-fold compared to the corresponding *scro^ΔE2-Gal4*^*/+ or *scro^ΔE3-Gal4*^*/+ controls.

We tested if the hybrid E2 (E2/EGFP) exon influences circRNA and/or lineRNA production. Since the knock-in mutants were homozygous lethal during late embryogenic and early larval stages, heterozygous flies were used to detect circE2/EGFP. RT-PCR results showed that splicing of intron-1 and intron-2 occurred normally to produce lineRNA containing E2/EGFP exon (Fig. 7A lower panel, lanes 2 and 3). We also detected single-exonic circRNA carrying E2/EGFP exon (Fig. 7A lower panel, lane 4); this was confirmed by sequencing (Supplementary Fig. 4). In addition, bi-exonic circE2/EGFP-E3 was produced from these heterozygous flies (Fig.7A lower panel, lane 5). These results are consistent with the previous finding that the exon sequence does not significantly contribute to the circularization process (Kramer *et al*. 2015).

To verify the requirement of splicing junction sites for the circRNA biogenesis, we took advantage of the *scro^ΔE3-EGFP*^* line in which EGFP knock-in deletes the 5’ portion of intron-3 including the 5’ splicing site (Fig. 7B). Accordingly, we anticipated the lack of splicing of the intron-3. RT-PCR results confirmed that the intron-2 is spliced out, but the intron-3 is not spliced out and remains in the linear transcript derived from *scro^ΔE3-EGFP*^*(Fig. 7B lower panel, lanes 2 and 3). The linear transcript containing the intron-3 was not shown in Fig. 7B, because of the short extension time for PCR conditions. Meanwhile, three different primer sets intended to detect circRNAs derived from the E3/EGFP exon did not show any products (Fig. 7B lower panel, lanes 4-6), supporting that the splice site is required for circularization. Based on these observations, we conclude that the circularization process does not need the endogenous exon sequences but the exon-intron splice site.

In our curiosity, we also wanted to see if overexpressed Scro could alter the production levels of *scro*’s lineRNA and/or circRNAs. To test it, Gal4 knock-in lines were crossed to *UAS-scro^HA^* (Nair *et al*. 2020). Since the UAS-*scro^HA^*transgene does not contain E1 and any of the introns, none of the circRNAs are expected to be produced from this construct. We then measured levels of lineRNAs and circRNAs derived ‘exclusively’ from the endogenous gene by RT-qPCR. For the lineRNAs, a set of E1F2 and E2R2 primers was used as the *UAS-scro^HA^* transgene-derived mRNA lacks the E1; for the circE2, E2-2F and E2R2 primers were employed since the binding site for E2R2 primer is absent in the Gal4 transgene (Supplementary Fig. 5). Interestingly, we found that the lineRNA levels increased by 5-7-fold (Fig. 7Ca), inferring an auto-regulatory mechanism in which Scro enhances the expression of its gene. On the other hand, we found that circE2 levels were significantly decreased 3-fold than the control (Fig. 7Ca). Progeny from a crossing of *UAS-scro^HA^* and *scro^ΔE3-Gal4*^* showed similar results (Fig. 7Cb). These results are somewhat consistent with the foregoing data (Fig. 6B) in which lower levels of circRNA are accompanied by higher levels of lineRNA. These data imply that Scro somehow regulates the ratio between linear and circular RNA forms by a novel mechanism. We speculate that the balanced production of lineRNA and circRNA is subjected to change in response to various cellular environments that alter the production of one or the other RNA form.

## DISCUSSION

The circRNAs are first thought to have arisen from aberrant splicing of pre-mRNAs (Cocquerelle *et al*. 1992, 1993). However, it is now widely accepted that circRNAs are one of the major classes of non-coding RNAs, and their biogenesis is delicately regulated (Salzman *et al*. 2012; Kristensen *et al*. 2019). In *Drosophila*, more than 2500 circRNAs have been annotated (Ashwal-Fluss *et al*. 2014; Westholm *et al*. 2014). However, the biogenesis of most circRNAs is little understood. In this study, we undertook comprehensive molecular analyses of *scro*-derived circRNAs to gain insights into the biogenesis mechanism of *scro* circRNAs in both *in vivo* and cell-culture systems as well as their possible roles.

We identified 12 circRNAs including the one previously reported (Westholm *et al*. 2014). Interestingly, all of them are either single- or multi-exonic circRNAs. The presence of multi-exonic ones indicates that inter-exonic introns are processed out after or before the formation of circularization, supporting that conventional splicing for lineRNAs and back-splicing events might utilize the same or similar splicing mechanism(s). Interestingly six *scro* circRNAs start with E2 and four with E3; since E3 is the second exon of *RF* transcript, our results are consistent with previous reports showing the second-exon biased circularization and co-transcriptional splicing for circRNA formation (Ashwal-Fluss *et al*. 2014; Westholm *et al*. 2014; Zhang *et al*. 2014). We did not find circRNAs containing E1 or E7. It is consistent with previous findings showing that the first or last exon of a gene is rarely made into circRNAs because they are flanked by only one intron (Salzman *et al*. 2012; Lasda and Parker 2014; Westholm *et al*. 2014; Gruner *et al*. 2016). We also did not find circRNAs containing E6, although this exon is flanked by the two largest introns.

Back-splicing mediated circularization is expected to involve both *cis*- and *trans*-factors (Kramer *et al*. 2015; Kristensen *et al*. 2019). As for the *cis*-factors, we showed that a hairpin structure involving 105-bp ICS in the E2-flanking introns is crucial for circE2 formation both in cultured cells and *in vivo*. In addition to the ICS, the regions immediately upstream and downstream of E2 are likely to be important (Figs. 4 and 7). Since these regions contain splicing junction sites, it seems that they are required for the basal level of back-splicing whereas ICS enhances this event significantly.

Another ICS pair located in intron-2 and intron-4 is likely to play a role in enhancing circE3-E4 production. Further aligning sequences of introns flanking other exons did not reveal candidate ICSs. The presence of ICSs is accountable for the circE2 and circE3-E4 being the two most abundant forms. Another notable feature of the ICSs is that the distance between the ICS and the splicing junction site influences the circularization efficiency. As shown (Fig. 4A), the ICS closer to the splicing junction site increased circRNA production levels perhaps via enhancing the hairpin formation.

ICSs mediate the formation of a hairpin structure that divides into two distinct parts, a loop and a stem. Our results showed that the most distal region from the loop (d-e in Fig. 5A) is critical for circRNA production, although the degree of base-pairing is greater in the proximal a-b region (Supplementary Fig. 2). Perhaps the distal region is either important for initiating the ICS pairing or for binding a *trans*-acting factor that is involved in the circularization process.

Recent studies showed that the back-splicing also needs *trans*-factors, and currently, a few RNA-binding proteins (RBP) are found to play roles in the biogenesis of *Drosophila* circRNAs (Kramer *et al*. 2015; Pamudurti *et al*. 2022). For example, splicing factors, hnRNP (heterogenous nuclear ribonucleoprotein), and SR (serine-arginine) proteins combinatorially regulate the circRNA levels derived from the *Drosophila laccase2* and *Plexin A,* but not those from *mbl* (Kramer *et al*. 2015). These studies suggest a mechanism in which sequence-specific RBPs regulate circRNA biogenesis in a gene-specific manner. Interestingly, transgenic over-expression of *scro^HA^* resulted in a significant reduction in circRNA production but an increase in lineRNAs from the endogenous locus (Fig. 7C). Since Scro functions as a homeodomain-transcription factor, it is unlikely that overexpressed Scro^HA^ suppresses circularization directly (Nair *et al*. 2020; Yoo *et al*. 2020; Konstantinides *et al*. 2022). Therefore, a conceivable idea is that Scro^HA^ directly regulates production levels of lineRNAs, which in turn results in the suppression of circRNA production. Alternatively, it could be an indirect effect through the expression of a Scro-target gene that encodes either an RBP or an RBP-regulating factor, which in turn controls the ratio of linear RNA to circRNA. In this case, identifying RBPs binding to the *scro*-ICSs and flanking intron sequences will shed light on the biogenesis mechanism of *scro*-circRNAs.

The molecular functions of circRNA are beginning to be understood (reviewed by Li *et al*. 2018; Kristensen *et al*. 2019). Relatively well-known roles of circRNAs include the regulation of gene expression via their binding to microRNAs (miRNA sponge) or RBPs (protein sponge). Searching for the miRBase (Kozomara and Griffiths-Jones, 2011) revealed miR-958-3p and miR-994-5p that show a substantial degree of complementarity to *scro* exon-2 and exon-4, respectively (Supplementary Fig. 6). While the function and biological process of miR-994 are unknown, miR-958 was reported to regulate Hedgehog signaling-mediated patterning of the wing imaginal disc (He *et al*. 2022). It will be interesting to investigate if *scro* circRNAs play any role as a sponge of these miRNAs, thereby affecting expression levels of these miRNA-target genes. A few exonic circRNAs are translated to produce truncated proteins which may or may not be functional (e.g., Pamudurti *et al*. 2017). We also found that certain *scro* circRNAs contain short open-reading frames that potentially produce truncated peptides. The roles of these peptides, if they are truly produced, will be an interesting subject of future study.

Biological roles of circRNAs may be associated with aging. We have shown that the levels of circE2 progressively escalated in aged fly heads. Likewise, the aging-dependent accumulation of circRNAs in fly heads is also observed for *mbl*, *CaMK-like*, *p120 catenin*, and *ankyrin 2* genes (Ashwal-Fluss *et al*. 2014; Westholm *et al*. 2014). Hence, circRNAs are considered a new indicator of the aging brain in *Drosophila* and mammals (Westholm *et al*. 2014; Gruner *et al*. 2016; Knupp and Miura 2018). Interestingly, aging *Drosophila* eyes show altered expression of splicing-related genes, which are causally related to an accumulation of circRNAs and visual senescence (Hall *et al*. 2017; Stegeman *et al*. 2018). Since *scro* transcripts are enriched in the optic lobe medulla in adult flies (Yoo *et al*. 2020), it will be interesting to resolve if accumulated *scro*-circRNA in the aged visual nervous system is also due to altered splicing factors and whether or not it contributes to aging-associated decline of visual function.

## Data availability

The authors affirm that all data necessary for confirming the conclusions of the article are present within the article, figures, and tables. Stocks are available upon request.

Supplemental material available at G3 online.

## Acknowledgments

We thank the Korea *Drosophila* Resource Center (KDRC) for the generation of deletion lines. This research is supported by the National Research Foundation of Korea (NRF) grant 2020R1I1A3074467 to SY and in part by an NIH grant (R15-GM140423) to JP.

## Conflicts of interest

The authors declare no conflicts of interest.

## SUPPLEMENTAL MATERIALS

**Supplementary Table 1.**
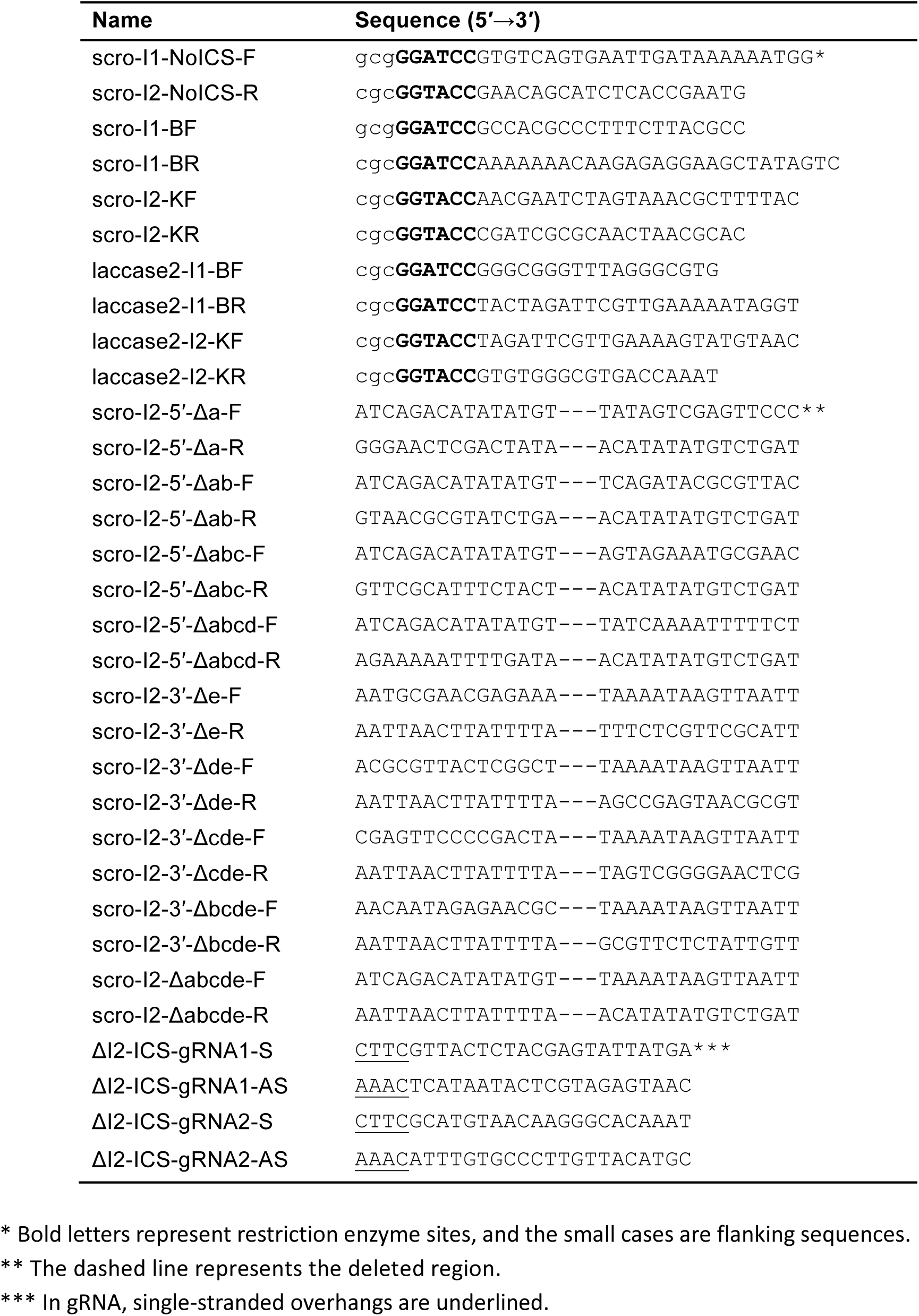
List of primers used for cloning.

**Supplementary Table 2.**
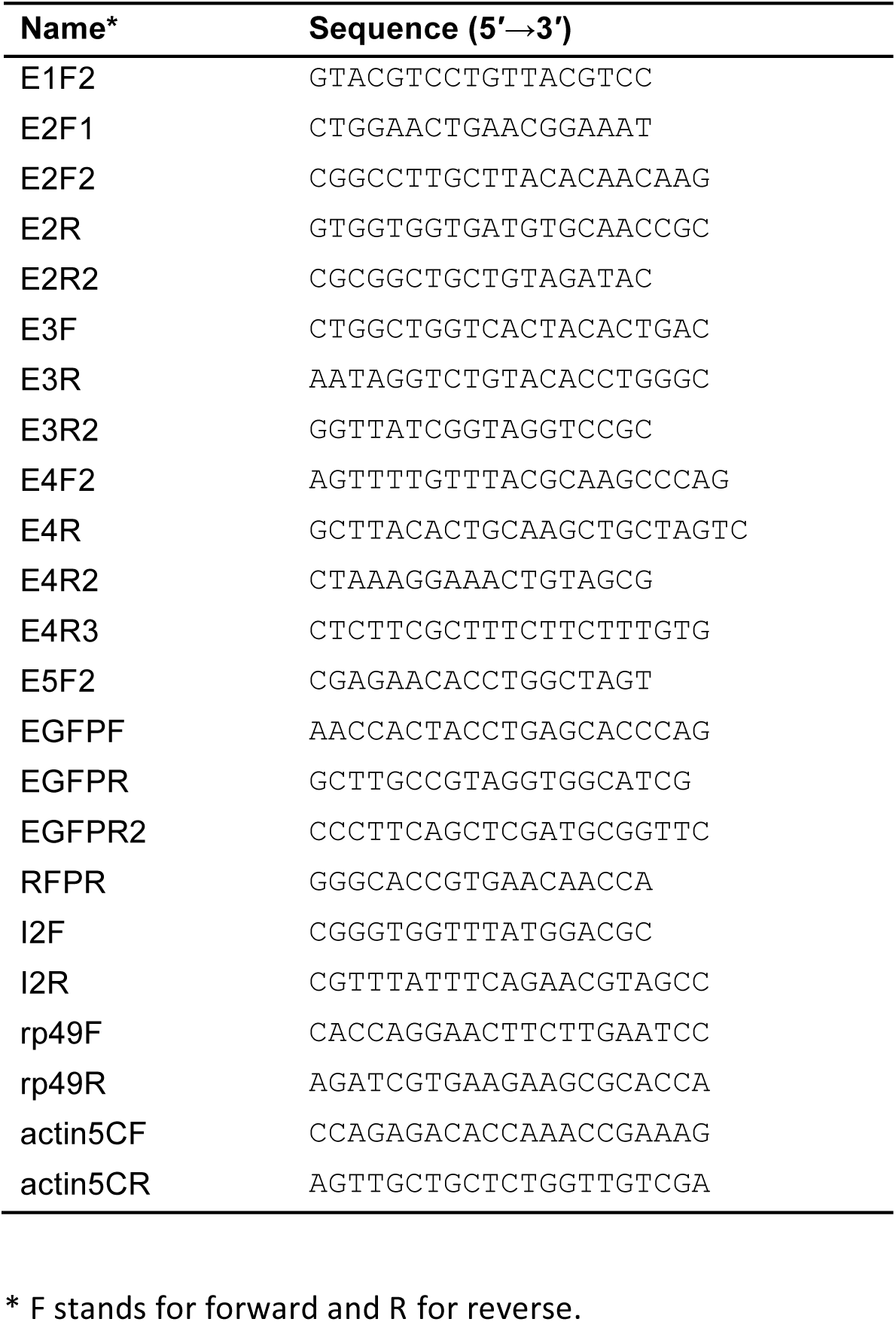
List of primers used for PCR.

**Supplementary Table 3.**
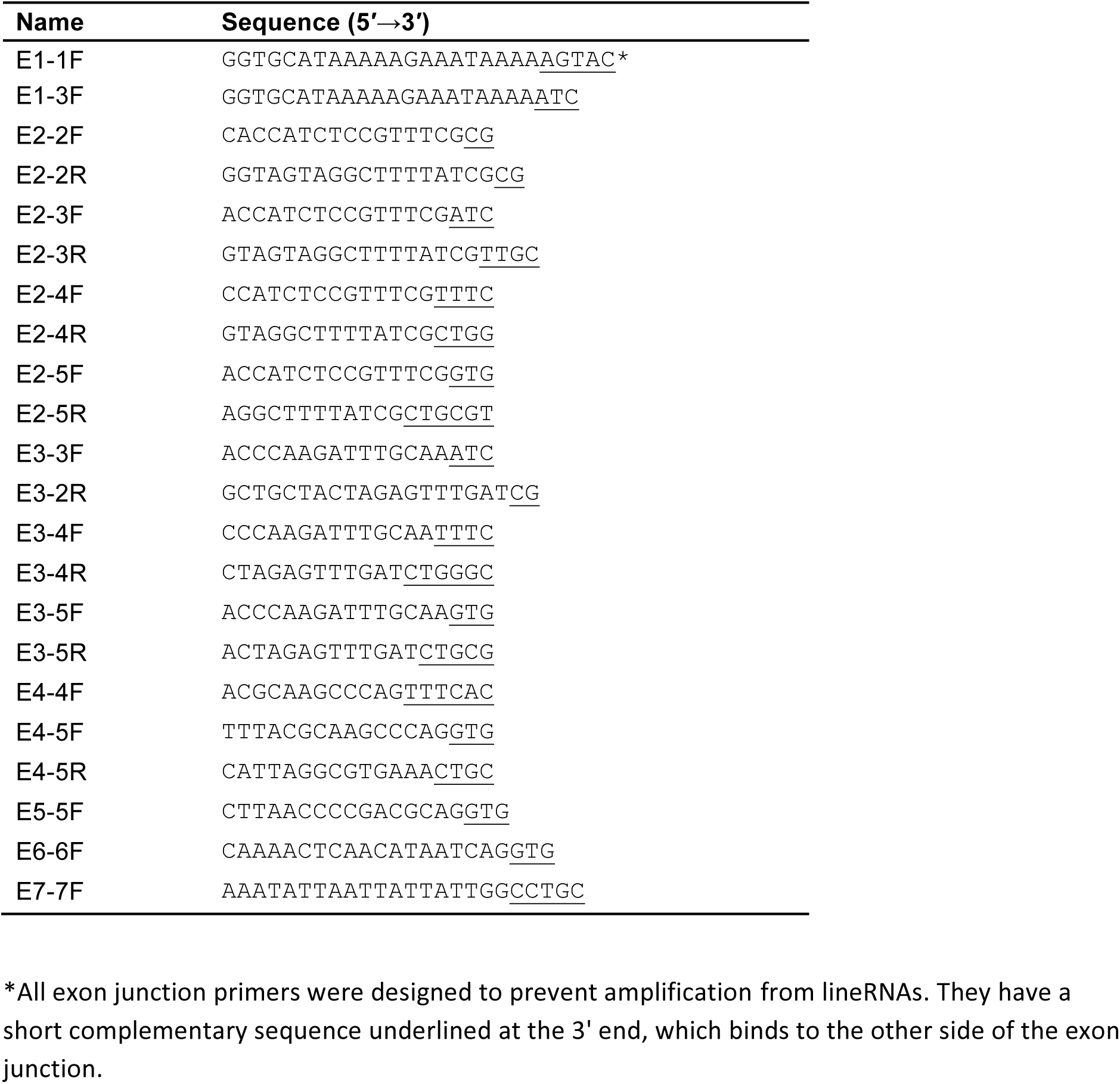
List of exon junction primers used for PCR.

**Supplementary Fig. 1.**
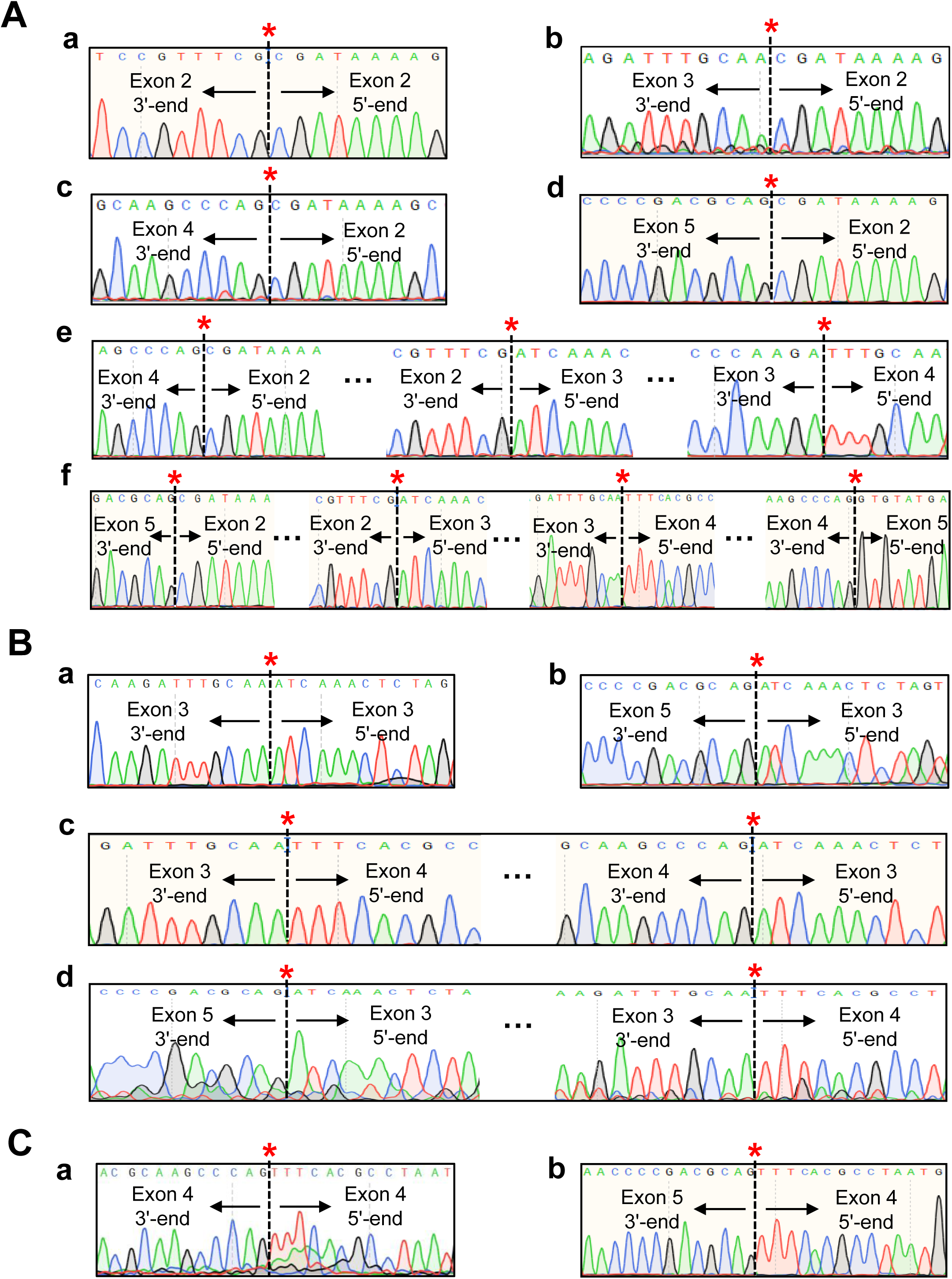
Sequencing data showing exon junction of various circRNAs. To identify all circRNAs derived from *scro* transcripts, cDNAs were synthesized using gene-specific primers, and PCR was carried out using either exon junction or divergent primers. The red asterisks (*) indicate the exon-exon joining site as a result of back-splicing. **(A)** Head-to-tail splice junctions of circRNAs containing exon-2 (E2) are derived from single-exon circular form, circE2 **(a)**, bi-exonic circE2-E3, circE2-E4, and cirE2-E5 **(b-d)**, tri-exonic circE2-E3-E4 **(e)**, and tetra-exonic circE2-E3-E4-E5 **(f)**. PCR primer sets and sequencing primers are as follows; circE2, E2-2F/E2R2 and E2R2; circE2-E3, E2-3F/E2-3R and E3F; circE2-E4, E2-4F/E2-4R and E4F2; circE2-E5, E2-5F/E2-5R and E5F2; circE2-E3-E4, E2-3F/E2-4R and E4F2; circE2-E3-E4-E5, E2-3F/E2-5R and E5F2, respectively. **(B)** Head-to-tail splice junctions of circRNAs containing exon-3 (E3) are derived from single-exon circular form, circE3 **(a)**, bi-exonic circE3-E5 and circE3-E4 **(b-c)**, and tri-exonic circE3-E4-E5 **(d)**. PCR primer sets and sequencing primers are as follows; circE3, E3-3F/E3R and E3R; circE3-E4, E3-4F/E3-4R and E4F2; circE3-E5, E3-5F/E3-5R and E5F2; circE3-E4-E5, E3-4F/E3-5R and E5F2, respectively. **(C)** Head-to-tail splice junctions of circRNAs containing exon-4 (E4) are derived from single-exonic circE4 **(a)** and bi-exonic circE4-E5 **(b)**. PCR primer sets and sequencing primers are as follows; circE4, E4-4F/E4R2 and E4R; circE4-E5, E4-5F/E4R and E5F2, respectively.

**Supplementary Fig. 2.**
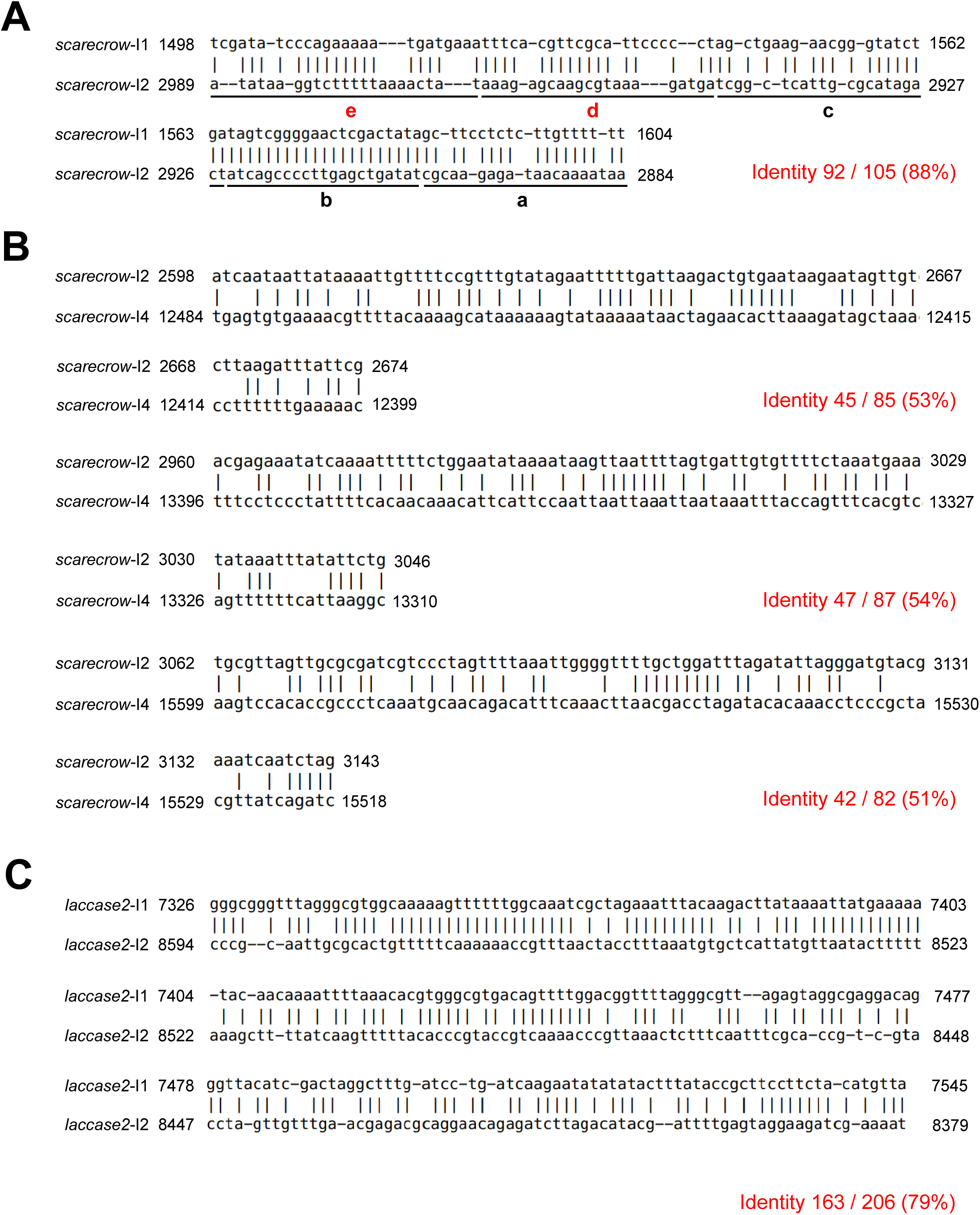
Identification of intronic complementary sequences (ICSs) in introns. To find candidate ICSs required for the production of circRNAs, intron sequences were subjected to EMBOSS explorer einverted program (http://emboss.toulouse.inra.fr/cgi-bin/emboss/einverted). The numbers indicate positions relative to the transcription start site. **(A)** Base-pairing region between intron-1 (I1) and intron-2 (I2) shows 105-bp ICS for *scro* circE2 (Figs. 4-6). **(B)** Three candidate ICS regions formed by base-pairing between intron-2 (I2) and intron-4 (I4) for *scro* circE3-E4 (blue boxes in Fig. 6A). **(C)** Base-pairing region between intron-1 (I1) and intron-2 (I2) from the *laccase2* gene displays 206-bp ICS for *laccase2* circRNA (Fig. 4Ac).

**Supplementary Fig. 3.**
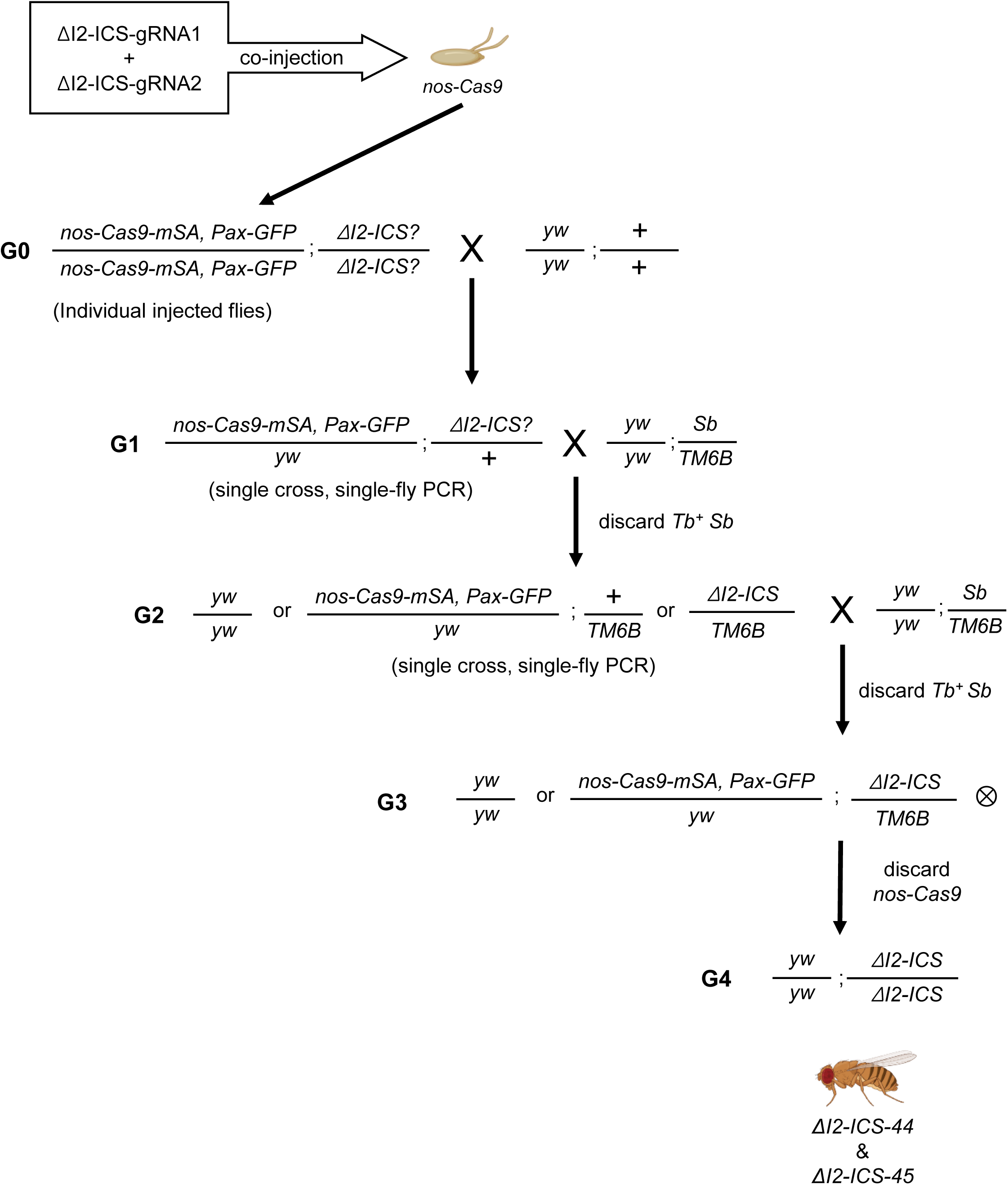
Generation of the *scro* mutants lacking ICS. *nos-Cas9* embryos were injected with a mixture of both ΔI2-ICS-gRNA1 and ΔI2-ICS-gRNA2 expression plasmids. Fifty G0 individuals were singly crossed to the *y w*. For each G0 line, 10 G1 offspring were collected and singly crossed with *y w*; *Sb/TM6B*. After 4 days, each G1 was subjected to single-fly PCR to identify the deletion. After the selection of positive G1 lines, about 5 to 6 G2 individuals were singly crossed with *y w*; *Sb/TM6B*, and single-fly PCR was performed again to verify the deletion of the target region. The G3 progeny was self-crossed to generate Δ*I2-ICS* homozygotes (G4). Fly images were created on BioRender (https://www.biorender.com/).

**Supplementary Fig. 4.**
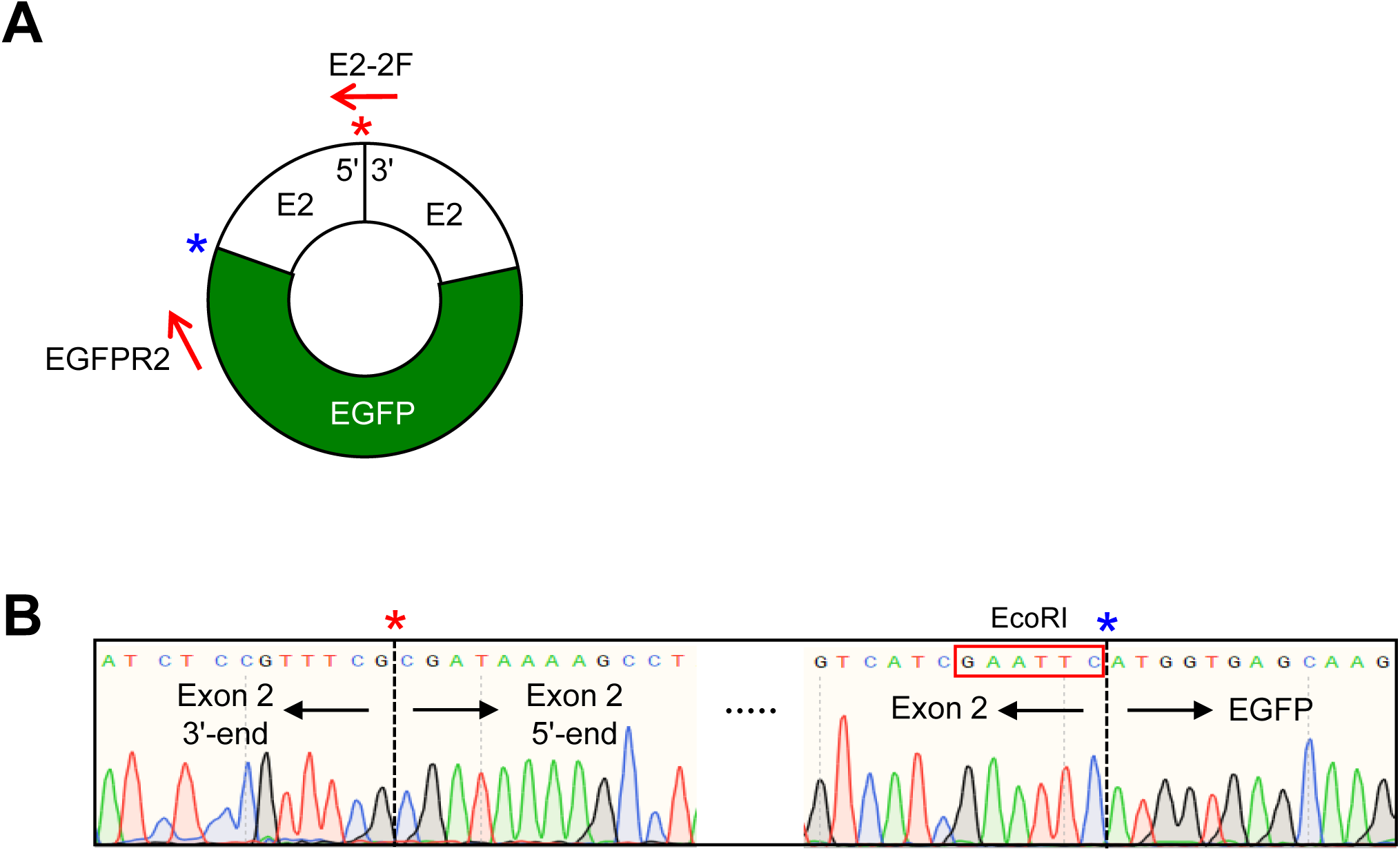
Verification of circRNA carrying E2/EGFP hybrid exon. **(A)** Schematic diagram of circRNA generated from *scro^ΔE2-EGFP*^* knock-in mutant which replaces part of exon-2 with EGFP-coding sequence. **(B)** Sequencing data showing head-to-tail splice junction of E2 (red asterisk) and E2/EGFP junction site (blue asterisk). The red box indicates the *EcoR*I site which is used for cloning to generate knock-in mutant flies.

**Supplementary Fig. 5.**
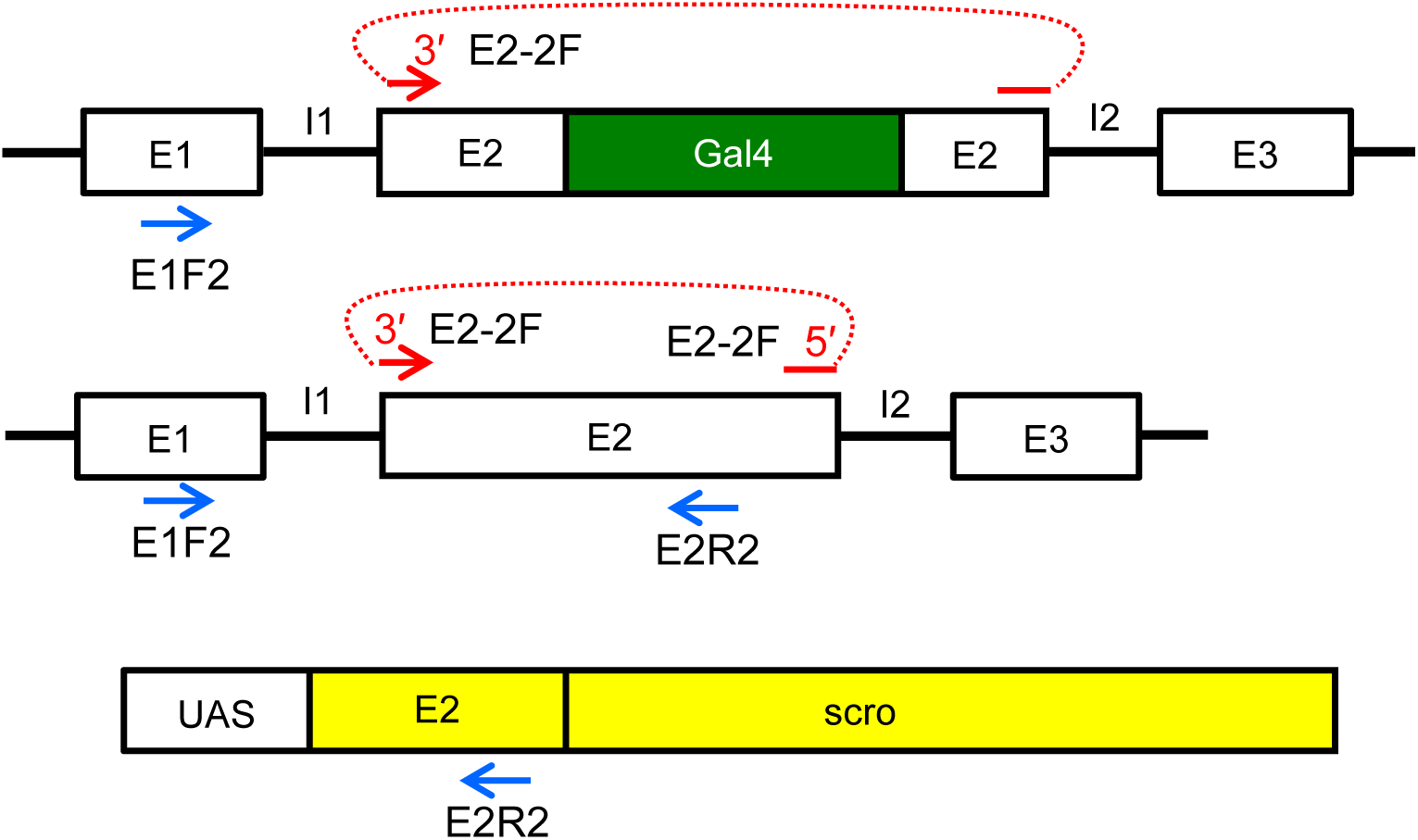
PCR strategy for detecting lineRNA and circE2 in Gal4 knock-in flies. PCR strategy related to Fig. 7Ca of detecting *scro* lineRNA and circE2 exclusively from the native exon; due to the lack of E2R2 binding site in the Gal4 coding sequence, circE2 is from native E2, not from Gal4 knock-in one (upper panel). Lack of E1 in the *UAS-scro* transgene disallows any PCR product (lower panel).

**Supplementary Fig. 6.**
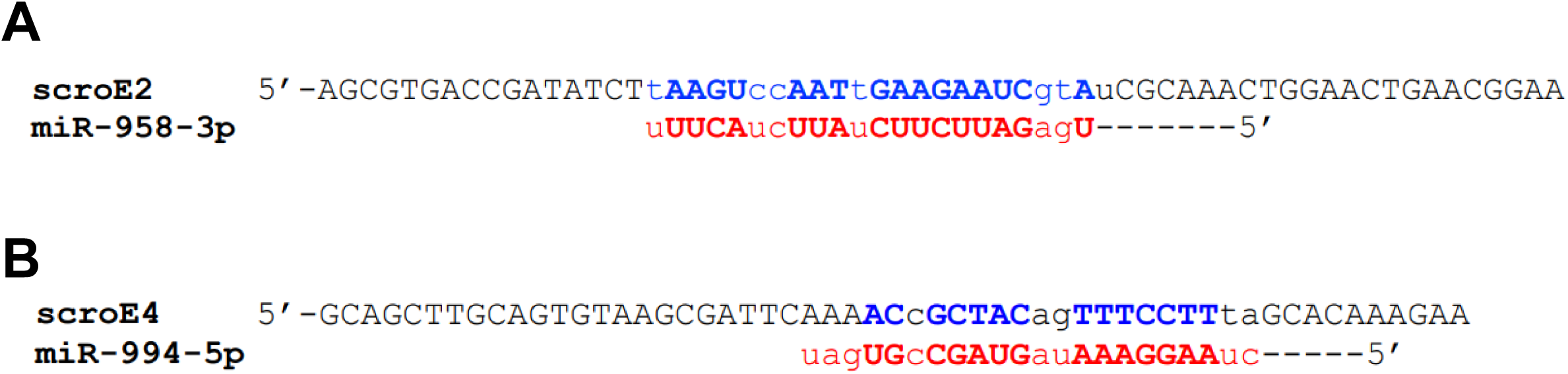
Candidate miRNAs targeted by *scro* circRNA. By using the miRBase web tool (https://www.mirbase.org/), we found candidate potential target sites of miRNA that can bind to *scro*-circRNA. **(A)** Complementarity between a region of *scro* E2 (blue) and miR-958-3p. **(B)** Complementarity between a region of *scro* E4 and miR-994-5p. Pairing nucleotides are shown in capitals.

## Literature cited

Aktaş T, Avşar Ilık İ, Maticzka D, Bhardwaj V, Pessoa Rodrigues C, Mittler G, Manke T, Backofen R, Akhtar A. 2017. DHX9 suppresses RNA processing defects originating from the ALU invasion of the human genome. Nature 544(7648):115–9. DOI: 10.1038/nature21715

Ashwal-Fluss R, Meyer M, Pamudurti NR, Ivanov A, Bartok O, Hanan M, Evantal N, Memczak S, Rajewsky N, Kadener S. 2014. circRNA biogenesis competes with pre-mRNA splicing. Mol Cell 56(1):55–66. DOI: 10.1016/j.molcel.2014.08.019

Barrett SP and Salzman J. 2016. Circular RNAs: Analysis, expression and potential functions. Development (Cambridge) 143(11):1838–47. DOI: 10.1242/dev.128074

Cha-Aim K, Hoshida H, Fukunaga T, Akada R. 2012. Fusion PCR via novel overlap sequences. Methods Mol Biol 852:97–110. DOI: 10.1007/978-1-61779-564-0_8

Cocquerelle C, Daubersies P, Majérus MA, Kerckaert JP, Bailleul B. 1992. Splicing with inverted order of exons occurs proximal to large introns. Embo J 11(3):1095–8. DOI: 10.1002/j.1460-2075.1992.tb05148.x

Cocquerelle C, Mascrez B, Hétuin D, Bailleul B. 1993. Mis-splicing yields circular RNA molecules. FASEB J 7(1):155–60. DOI: 10.1096/fasebj.7.1.7678559

Dubin RA, Kazmi MA, Ostrer H. 1995. Inverted repeats are necessary for circularization of the mouse testis sry transcript. Gene 167(1):245–8. DOI: 10.1016/0378-1119(95)00639-7

Gratz SJ, Ukken FP, Rubinstein CD, Thiede G, Donohue LK, Cummings AM, O’Connor-Giles KM. 2014. Highly specific and efficient CRISPR/Cas9-catalyzed homology-directed repair in *Drosophila*. Genetics 196(4):961–71. DOI: 10.1534/genetics.113.160713

Gruner H, Cortés-López M, Cooper DA, Bauer M, Miura P. 2016. CircRNA accumulation in the aging mouse brain. Sci Reports 6(1):38907. DOI: 10.1038/srep38907

Hansen T, Jensen T, Clausen B. 2013. Natural RNA circles function as efficient microRNA sponges. Nature 495:384–388. DOI: 10.1038/nature11993

Hall H, Medina P, Cooper DA, Escobedo SE, Rounds J, Brennan KJ, Vincent C, Miura P, Doerge R, Weake VM. 2017. Transcriptome profiling of aging *Drosophila* photoreceptors reveals gene expression trends that correlate with visual senescence. BMC Genomics 18(1):894. DOI: 10.1186/s12864-017-4304-3

He T, Fan Y, Wang Y, Liu M, Zhu AJ. 2022. Dissection of the microRNA Network Regulating Hedgehog Signaling in *Drosophila*. Front. Cell Dev Biol 10:866491. DOI: 10.3389/fcell.2022.866491

Ivanov A, Memczak S, Wyler E, Torti F, Porath HT, Orejuela MR, Piechotta M, Levanon EY, Landthaler M, Dieterich C, et al. 2015. Analysis of intron sequences reveals hallmarks of circular RNA biogenesis in animals. Cell Reports 10(2):170–7. DOI: 10.1016/j.celrep.2014.12.019

Jeck WR, Sorrentino JA, Wang K, Slevin MK, Burd CE, Liu J, Marzluff WF, Sharpless NE. 2013. Circular RNAs are abundant, conserved, and associated with ALU repeats. RNA (Cambridge) 19(2):141–57. DOI: 10.1261/rna.035667.112

Jeck WR and Sharpless NE. 2014. Detecting and characterizing circular RNAs. Nat Biotechnol 32(5):453–61. DOI: 10.1038/nbt.2890

Knupp D and Miura P. 2018. CircRNA accumulation: A new hallmark of aging? Mech Ageing Dev 173:71–9. DOI: 10.1016/j.mad.2018.05.001

Knupp D, Cooper DA, Saito Y, Darnell RB, Miura P. 2021. NOVA2 regulates neural circRNA biogenesis. Nuc Acids Res 49(12):6849–62. DOI: 10.1093/nar/gkab523

Kozomara A, Griffiths-Jones S. 2011. miRBase: integrating microRNA annotation and deep-sequencing data. Nuc Acids Res 39(Database issue): D152–D157. DOI: 10.1093/nar/gkq1027

Konstantinides N, Holguera I, Rossi AM, Escobar A, Dudragne L, Chen YC, Tran TN, Martínez Jaimes AM, Özel MN, Simon F, et al. 2022. A complete temporal transcription factor series in the fly visual system. Nature 604(7905): 316–322. DOI: 10.1038/s41586-022-04564-w

Kramer MC, Liang D, Tatomer DC, Gold B, March ZM, Cherry S, Wilusz JE. 2015. Combinatorial control of *Drosophila* circular RNA expression by intronic repeats, hnRNPs, and SR proteins. Genes Dev 29(20):2168–82. DOI: 10.1101/gad.270421.115

Kristensen LS, Andersen MS, Stagsted LVW, Ebbesen KK, Hansen TB, Kjems J. 2019. The biogenesis, biology, and characterization of circular RNAs. Nat Rev Genet 20(11):675–91. DOI: 10.1038/s41576-019-0158-7

Lasda E and Parker R. 2014. Circular RNAs: Diversity of form and function. RNA (Cambridge) 20(12):1829–42. DOI: 10.1261/rna.047126.114

Li X, Yang L, Chen L. 2018. The biogenesis, functions, and challenges of circular RNAs. Mol Cell 71(3):428–42. DOI: 10.1016/j.molcel.2018.06.034

Liang D and Wilusz JE. 2014. Short intronic repeat sequences facilitate circular RNA production. Genes Dev 28(20):2233–47. DOI: 10.1101/gad.251926.114

Memczak S, Jens M, Elefsinioti A, Torti F, Krueger J, Rybak A, Maier L, Mackowiak SD, Gregersen LH, Munschauer M, et al. 2013. Circular RNAs are a large class of animal RNAs with regulatory potency. Nature 495(7441):333–8. DOI: 10.1038/nature11928

Nair S, Bahn JH, Lee G, Yoo S, Park JH. 2020. A homeobox transcription factor scarecrow (SCRO) negatively regulates pdf neuropeptide expression through binding an identified cis-acting element in *Drosophila melanogaster*. Mol Neurobiol 57(4):2115–30. DOI: 10.1007/s12035-020-01874-w

Pamudurti NR, Patop IL, Krishnamoorthy A, Bartok O, Maya R, Lerner N, Ashwall-Fluss R, Konakondla JVV, Beatus T, Kadener S. 2022. circMbl functions *in cis* and *in trans* to regulate gene expression and physiology in a tissue-specific fashion. Cell Reports (Cambridge) 39(4):110740. DOI: 10.1016/j.celrep.2022.110740

Pamudurti NR, Bartok O, Jens M, Ashwall-Fluss R, Stottmeister C, Ruhe L, Hanan M, Wyler E, Perez-Hernandez D, Ramberger E, Shenzis S, Samson M, Dittmar G, Landthaler M, Chekulaeva M, Rajewsky N, Kadener S. 2017. Translation of circRNAs. Mol Cell 66:9–21.e7 DOI: 10.1016/j.molcel.2017.02.021

Park JH, Helfrich-Förster C, Lee G, Liu L, Rosbash M, Hall JC. 2000. Differential regulation of circadian pacemaker output by separate clock genes in *Drosophila*. Proc Natl Acad Sci USA 97(7):3608–13. DOI: 10.1073/pnas.97.7.3608

Patop IL, Wüst S, Kadener S. 2019. Past, present, and future of circRNAs. EMBO J 38(16):e100836. DOI: 10.15252/embj.2018100836

Renn SC, Park JH, Rosbash M, Hall JC, Taghert PH. 1999. A pdf neuropeptide gene mutation and ablation of PDF neurons each cause severe abnormalities of behavioral circadian rhythms in *Drosophila*. Cell 99(7):791. DOI: 10.1016/s0092-8674(00)81676-1

Salzman J, Chen RE, Olsen MN, Wang PL, Brown PO. 2013. Cell-type specific features of circular RNA expression. PLOS Genetics 9(9):e1003777. DOI: 10.1371/journal.pgen.1003777

Salzman J, Gawad C, Wang PL, Lacayo N, Brown PO. 2012. Circular RNAs are the predominant transcript isoform from hundreds of human genes in diverse cell types. Plos One 7(2):e30733. DOI: 10.1371/journal.pone.0030733

Sanger HL, Klotz G, Riesner D, Gross HJ, Kleinschmidt AK. 1976. Viroids are single-stranded covalently closed circular RNA molecules existing as highly base-paired rod-like structures. Proc Natl Acad Sci USA 73(11):3852–6. DOI: 10.1073/pnas.73.11.3852

Shen T, Han M, Wei G, Ni T. 2015. An intriguing RNA species-perspectives of circularized RNA. Protein Cell 6(12):871–80. DOI: 10.1007/s13238-015-0202-0

Starke S, Jost I, Rossbach O, Schneider T, Schreiner S, Hung L, Bindereif A. 2015. Exon circularization requires canonical splice signals. Cell Reports 10(1):103–11. DOI: 10.1016/j.celrep.2014.12.002

Stegeman R, Hall H, Escobedo SE, Chang HC, Weake VM. 2018. Proper splicing contributes to visual function in the aging *Drosophila* eye. Aging Cell 17(5):e12817. DOI: 10.1111/acel.12817

Tang JLY, Hakes AE, Krautz R, Suzuki T, Contreras EG, Fox PM, Brand AH. 2022. NanoDam identifies Homeobrain (ARX) and Scarecrow (NKX2.1) as conserved temporal factors in the *Drosophila* central brain and visual system. Dev. Cell 57(9):1193–1207.e7. DOI: 10.1016/j.devcel.2022.04.008

Wang M, Hou J, Müller-McNicoll M, Chen W, Schuman EM. 2019. Long and repeat-rich intronic sequences favor circular RNA formation under conditions of reduced spliceosome activity. iScience 20:237–47. DOI: 10.1016/j.isci.2019.08.058

Wang PL, Bao Y, Yee M, Barrett SP, Hogan GJ, Olsen MN, Dinneny JR, Brown PO, Salzman J. 2014. Circular RNA is expressed across the eukaryotic tree of life. Plos One 9(3):e90859. DOI: 10.1371/journal.pone.0090859

Wang W, Liu W, Wang Y, Zhou L, Tang X, Luo H. 2011. Notch signaling regulates neuroepithelial stem cell maintenance and neuroblast formation in *Drosophila* optic lobe development. Dev Biol 350(2):414–28. DOI: 10.1016/j.ydbio.2010.12.002

Weigelt CM, Sehgal R, Tain LS, Cheng J, Eßer J, Pahl A, Dieterich C, Grönke S, Partridge L. 2020. An insulin-sensitive circular RNA that regulates lifespan in *Drosophila*. Mol Cell 79(2):268,279.e5. DOI: 10.1016/j.molcel.2020.06.011

Westholm JO, Miura P, Olson S, Shenker S, Joseph B, Sanfilippo P, Celniker SE, Graveley BR, Lai EC. 2014. Genome-wide analysis of *Drosophila* circular RNAs reveals their structural and sequence properties and age-dependent neural accumulation. Cell Reports 9(5):1966–80. DOI: 10.1016/j.celrep.2014.10.062

Yoo S, Nair S, Kim H, Kim Y, Lee C, Lee G, Park JH. 2020. Knock-in mutations of scarecrow, a *Drosophila* homolog of mammalian Nkx2.1, reveal a novel function required for development of the optic lobe in *Drosophila melanogaster*. Dev Biol 461(2):145–59. DOI: 10.1016/j.ydbio.2020.02.008

Zaffran S, Das G, Frasch M. 2000. The NK-2 homeobox gene scarecrow (scro) is expressed in pharynx, ventral nerve cord, and brain of *Drosophila* embryos. Mech Dev 94(1):237–41. DOI: 10.1016/s0925-4773(00)00298-7

Zhang X, Wang H, Zhang Y, Lu X, Chen L, Yang L. 2014. Complementary sequence-mediated exon circularization. Cell 159(1):134–47. DOI: 10.1016/j.cell.2014.09.001

Zheng Q, Bao C, Guo W, Li S, Chen J, Chen B, Luo Y, Lyu D, Li Y, Shi G, et al. 2016. Circular RNA profiling reveals an abundant circHIPK3 that regulates cell growth by sponging multiple miRNAs. Nat Commun 7(1). DOI: 10.1038/ncomms11215

